# Dichloroacetate improves animal survival, growth, neuromuscular activity, mitochondrial stress and physiology, and elevated lactate in *C. elegans pdha-1* and *dld-1* RNAi models of pyruvate dehydrogenase complex deficiency (PDCD)

**DOI:** 10.64898/2026.07.08.737008

**Authors:** Cristina Remes, Neal D. Mathew, Victoria Miranda, Suraiya Haroon, Tyler O’Hara, Vernon E. Anderson, Manuela Lavorato, Kelsey Keith, Rui Xiao, Eiko Nakamaru-Ogiso, Marni J. Falk

**Affiliations:** Mitochondrial Medicine Frontier Program, Division of Human Genetics, Department of Pediatrics, The Children’s Hospital of Philadelphia (CHOP), Philadelphia, Pennsylvania, USA; Department of Pediatrics, Perelman School of Medicine, University of Pennsylvania, Philadelphia, Pennsylvania, USA; Department of Biomedical and Health Informatics, The Children’s Hospital of Philadelphia, Philadelphia, PA; Department of Biostatistics, Perelman School of Medicine, University of Pennsylvania, Philadelphia

**Keywords:** Mitochondria, growth, neuromuscular activity, pyruvate dehydrogenase complex, *PDHA1*, *DLD*, RNA interference, dichloroacetate (DCA)

## Abstract

Pyruvate dehydrogenase complex (PDHc) deficiency (PDCD) is a primary mitochondrial disorder characterized by neurodevelopmental disability, altered intermediary metabolism and early mortality. Dichloroacetate (DCA), a pyruvate analogue, is a well-described PDHc activator that remains under clinical investigation for treatment of PDCD. Here, we studied the *in vivo* efficacy of a 5-point log concentration range of DCA on animal health and metabolism in *C. elegans* with feeding RNA interference (RNAi) expression knockdown of either *PDHA-1* or *DLD-1* homologues at graded degrees to model variable disease severity. These worm models recapitulate phenotypic features of PDCD observed in human patients, including reduced survival, delayed growth, locomotor impairment, and elevated lactate and/or pyruvate tissue levels. DCA treatment appeared well-tolerated, with no gross morphologic toxicity seen at doses up to 25 mM. Significantly improved health, survival, tissue lactate levels, and mitochondrial physiology were observed at 25 mM in *pdha-1(RNAi)* knockdown animals. DCA treatment in *dld-1(RNAi) C. elegans* models (undiluted, 1:20 dilution, and 1:100 dilution) showed significant therapeutic benefits on survival, neuromuscular function and metabolic phenotypes primarily in the moderate (1:20) and/or mild (1:100) *dld-1(RNAi)* deficiency strains, but not in full-dose *dld-1(RNAi)*. Importantly, linear growth, neuromuscular activity, and mitochondrial physiology were significantly improved with DCA treatment even in the most severe *dld-1(RNAi)* undiluted model. Overall, preclinical modeling provides objective evidence of DCA therapeutic efficacy in *C. elegans* expression knockdown strains for two well-conserved homologues of *PDHA1* and *DLD* that represent distinct genetic etiologies of PDHc deficiency, with demonstrated beneficial effects on survival, healthspan, tissue lactate, and mitochondrial physiology. These data further confirm that DCA’s therapeutic effect correlates with PDHc disease phenotype severity in *dld-1(RNAi)* animals.

**SYNOPSIS: Dichloroacetate (DCA) treatment demonstrated significant preclinical beneficial effect on survival, neuromuscular function, linear growth, tissue lactate, and mitochondrial metabolism in two *C. elegans* models with variable degrees of pyruvate dehydrogenase complex (PDHc) deficiency (PDCD), providing confirmatory evidence to support its therapeutic potential in human PDCD patients.**

## INTRODUCTION

Pyruvate dehydrogenase complex (PDHc) deficiency (PDCD, OMIM 312170) is a rare genetic mitochondrial disorder, mainly affecting the central nervous system with manifestations of neurodevelopmental delay, structural brain abnormalities, neuromuscular disability, lactic acidosis and increased plasma pyruvate^1-6^. Human PDHc is a large (approximately 9 megadalton), highly structured multi-subunit enzyme complex^7,8^ located in the mitochondrial matrix that catalyzes the irreversible oxidative decarboxylation of pyruvate into acetyl-CoA. Pathogenic variants in any of the genes encoding PDHc subunits can cause PDHc deficiency, where the most common occur in the gene encoding the pyruvate dehydrogenase E1α subunit (*PDHA1*, OMIM 300502) ^9^. Less commonly, pathogenic variants of dihydrolipoamide dehydrogenase, (*DLD,* OMIM 246900), cause 6% of PDHc deficiency cases^2^. In addition to its role in PDHc activity, DLD also serves as the dihydrolipoamide dehydrogenase (E3 subunit) in 4 other keto acid dehydrogenase complexes ^10-12^. Sequence alignment of human and *C. elegans* PDHA1 and DLD show a high degree of protein sequence homology, with 58.7% and 68.4% sequence identity and 96.7% and 96.9% sequence similarity, respectively (Supplementary Fig. 1A, Supplementary Fig. 7A).

Currently, no US Food and Drug Administration (FDA)-approved treatments or cures exist for primary PDHc deficiency (PDCD). However, beneficial effects of dichloroacetate (DCA) have been reported in patient case reports^13,14^, animal models^11^, and cellular models^15-18^. Two mechanisms by which DCA increases PDHc activity have been proposed. The first mechanism involves inhibition of PDHc kinases, thereby keeping PDHc in its unphosphorylated, catalytically active state^19,20^. Another possible mechanism that explains DCA beneficial effects in PDHc deficiency that has been explored in human fibroblasts is stabilization of the PDHc enzymatic complex to reduce its rate of degradation^15,21^. However, cellular models have been shown to have selective sensitivity to DCA^21^, the reason for which is currently not fully understood.

A multi-site, phase 3, randomized, placebo-controlled, cross-over clinical trial investigating the therapeutic effect of DCA in human PDCD subjects has been completed with regulatory review pending from the FDA ^22^. However, due to the inherent rare nature of this disease, it is not feasible to complete a second adequate and well controlled clinical trial of DCA effect in PDHc deficiency. Therefore, we sought to provide confirmatory, objective, pre-clinical evidence of dose-dependent safety and therapeutic efficacy of DCA as a treatment for PDCD. To this end, we used two *C. elegans* RNA interference (RNAi) knockdown strains (*pdha-1* and *dld-1*) to model PDCD ^11,23^ and evaluate the efficacy of DCA on animal survival, health, and diverse aspects of mitochondrial physiology. In addition, we used a graded RNAi feeding approach to characterize the *dld-1 (RNAi)* knockdown models at three different dilutions to evaluate DCA treatment effects in models with different degrees of DLD deficiency and phenotypic severity at the level of animal survival, linear growth, neuromuscular function, and intermediary metabolic function including tissue lactate and pyruvate levels and mitochondrial physiology assessments.

## 1. METHODS

An overview of the experimental design and conditions is provided in Supplementary Table 2.

### 1.1 *C. elegans* strains and maintenance

The original N2 Bristol, GL347 (*hsp-6p::GFP*), VS21 (*myo-2p::mCherry*), and the SJ4103 *C. elegans* strain, carrying the zcls14 construct (*myo-3p::GFP(mit)* were obtained from the *Caenorhabditis* Genetics Center (CGC) at the University of Minnesota. The GL347 (*hsp-6p::GFP*) and VS21 (*myo-2p::mCherry*) strains were crossed to generate a wild-type line expressing both *hsp-6p::GFP* and *myo-2p::mCherry*. Worms were maintained under standard conditions at 20°C on nematode growth media (NGM) seeded with *E. coli* OP50, unless stated otherwise. DCA was provided under sponsored research agreement from Saol Therapeutics, to evaluate the potential efficacy and tolerability of a human-grade formulation used in PDCD patient clinical trials, as opposed to prior study by this research group of commercially available laboratory grade DCA^11^.

### 1.2 Synchronization of *C. elegans*

Embryos were obtained from gravid adult animals by exposure to commercial concentrated germicidal bleach containing 8.25% sodium hypochlorite (Clorox). Worms were collected from NGM plates with 4 mL S. Basal (23.4 g NaCl, 4 g K_2_HPO_4_, 25 g KH_2_PO_4_, 4 mL cholesterol (5 mg/mL in ethanol), 4 L milliQ water, pH = 7). 1 mL fresh bleach and 500 µL 5N NaOH were added to the worm suspension and the tubes were mixed for approximately 2 min, until no visible worm bodies were present. The samples were centrifuged at 199*g* for 1 min, and the supernatant was immediately removed. The pellet was subsequently washed twice with 10 mL S. Basal followed by centrifugation at 1200*g* for 1 min.

### 1.3 RNAi Feeding knockdown of PDHA-1 and DLD-1 in *C. elegans*

Knockdown of PDHA-1 and DLD-1 in *C. elegans* was achieved by feeding transfected *E. coli*. *C. elegans* embryos were grown on NGM supplemented with 0.2 mM isopropyl β-D-1-thiogalactoside (IPTG), 100 µg/mL ampicillin, treated with DCA or water, and seeded with *pdha-1(RNAi)* (clone T05H10.6, *C. elegans* ORF Collection, Horizon), *dld-1(RNAi)* (clone LLC1.3, Ahringer RNAi collection, Source BioScience) or control RNAi (clone L4440, Ahringer RNAi collection, Source BioScience). Prior to the knockdown experiments, all clones were sequence verified. For most experiments the *dld-1(RNAi)* was undiluted or diluted with L4440 at 1:20 (*dld-1(1:20 RNAi)*) or 1:100 (*dld-1(1:100 RNAi)*). Unless stated otherwise, worms were analyzed at stage L4+1.

### 1.4 *C. elegans* gross morphology analysis

Wild-type (N2 Bristol) DLD-1 and PDHA-1 knockdown worms were grown on solid media treated or untreated with DCA and observed at L4 + 5 days of adulthood to visually identify toxicity at the level of gross morphologic toxicity and identify the maximal safe effective dosage exposure.

### 1.5 *C. elegans* lifespan analyses using WormScan

Wild-type (N2 Bristol), DLD-1 and PDHA-1 knockdown worms, treated with DCA or water were age-synchronized by bleaching as described above. Stage L4 animals were then transferred onto 6 cm plates (30-60 worms per plate) containing 50 µM 5-fluorodeoxyuridine (FUDR), 0.2 mM IPTG, 100 µM ampicillin and seeded with the corresponding RNAi bacteria, and treated with DCA or water. Two consecutive images (one image before and one image after light stimulation) were recorded every 48 h using an Epson V800 scanner. Worm survival was scored using WormScan Machine Learning (ML) v6.3, a Python desktop application developed in-house. The images are compared using two deep learning models. A YOLO26 object detector^24,25^ localized individual worms in each scanned image, and a MobileNetV2 classifier^26^ determined alive/dead status from a stitched side-by-side composite of the two timepoints. Each image pair was loaded into the application, the plate area was isolated using a circular crop, and any annotations or writing on the plate were masked. The YOLO26 detector (trained on 18 plates, 417 annotations, validation mAP50 = 0.91) identified worm locations in the T1 image. For each detection, a 256 × 128 pixel composite was generated by stitching the same region from both T0 and T1 timepoints, which was passed to the MobileNetV2 classifier (trained on 1425 alive and 251 dead examples; balanced validation accuracy = 98%) for alive/dead assignment. Detections with confidence below 0.25 (detector) or 0.50 (classifier) were excluded. Counts were exported for survival curve calculation, with manual correction available for any miscalled worms. The image analysis tool WormScan ML v6.3.1 (https://doi.org/10.5281/zenodo.21042471) is archived on Zenodo^27^.

### 1.6 Manual *C*. *elegans* lifespan analyses

Wild type (N2 Bristol), *pdha-1 (RNAi)* and *dld-1 (RNAi)* treated with 25 mM DCA or water were age-synchronized by bleaching. At the L4 stage, approximately 30 worms were transferred to 3.5 cm NGM plates supplemented with 0.2 mM IPTG and ampicillin 100 µg/mL, and seeded with the corresponding RNAi bacteria. Worms were scored daily for survival and transferred to fresh plates until reproduction ceased. Animals unresponsive to gentle stimulation with a wire pick were scored as dead. Animals lost due to non-natural causes were censored from survival analysis. Three biological replicates were analyzed for all conditions.

### 1.7 *C. elegans* light stimulated locomotor activity measurements

Automated locomotor activity analysis was conducted in 24-well plates prepared with transparent GelRite-based media to enable optimal imaging, using a previously described method ^28^. Each well contained 1 mL of media (8 g/L GelRite, 3 g/L NaCl, 2.5 g/L peptone, 40 µM FUDR, and 100 µM ampicillin) supplemented with either 0.2 mM or 2 mM IPTG for *dld-1* and *pdha-1* RNAi conditions, respectively. Media between wells consisted of 6.7 g/L GelRite and 3 g/L NaCl. Plates were treated with the appropriate concentration of DCA or water (buffer control), seeded with the corresponding RNAi bacteria and allowed to dry overnight before use. Wild-type (N2 Bristol), *dld-1*(*full-dose RNAi*), and *pdha-1(RNAi)* knockdown worms were age-synchronized by bleaching onto the appropriate 10 cm RNAi plates, as described above. L4 stage worms were collected with 10 mL S. Basal and concentrated to 2 worms/µL. 10 µL of worm suspension (20 worms) was added to each well and the plates were allowed to dry for 15 min. Six technical replicates were included per condition on each plate. Before imaging, plate lids were treated with anti-fog spray to prevent condensation. Plates were imaged twice daily, before and after blue-light stimulation, using the WormWatcher robot over a 30-day period, as previously described ^28^. Data w***ere*** analyzed and visualized using R ^29^.

### 1.8 *C. elegans* linear growth (length) measurements

Wild type (N2 Bristol), PDHA-1 and DLD-1 knockdown worms were imaged on solid media using an Epson V800 scanner, and the length of each individual worm was analyzed using ImageJ. Three biological replicates per condition were performed, with a total of 30 worms per condition. The lengths of the DLD-1 and PDHA-1 knockdown worms were normalized to the length of the wild-type worms.

### 1.9 Neuromuscular activity and locomotor behavior analyses in *C. elegans*

Motility of wild type (N2 Bristol), PDHA-1 and DLD-1 knockdown worms in liquid media was measured using a thrashing assay. Worms were placed in S. Basal and a 10 s video was collected using a Basler USB Camera (model #108014) with a KOWA industrial lens 75 mm/F2.5. The FIJI plugin wrMTrck ^30^ was used to determine the body bends/s of each individual *C. elegans*. Three biological replicate experiments were performed, with a total of 60 worms analyzed for each condition.

### 1.10 Mitochondrial unfolded protein response (UPR^mt^) induction analysis

The *C. elegans* N2 (*hsp-6p::gfp* + *myo-2p::mCherry*) strain was used to quantify induction of the UPR^mt31^. Wild-type, DLD-1 and PDHA-1 knockdown worms were grown from embryo stage on 10 cm NGM plates containing 0.2 mM IPTG, 100 µg/mL ampicillin, treated with DCA or buffer control and seeded with the corresponding RNAi bacteria. Day 1 adult worms were washed in S. Basal to a density of 1 worm/µL. The worm suspension was then dispensed into a 384 well-plate (384 Well Black Plate, Optically Clear Polymer Bottom, USA Scientific #5678-1091), at a volume of 50 µL/well. Subsequently, 50 µL of NaN_3_ (0.1 M supplemented with 0.01% Tween-20) was added to each well and incubated for 15 min at room temperature. Mitochondrial stress was then measured using a CellInsight CX5 High Content Screening system (ThermoFisher Scientific). The red channel was used to identify and quantify the number of worms in each well, and the total fluorescence detected in the green channel was used to quantify the total UPR^mt^ of each well. The total fluorescence measured in the green channel was then normalized to the number of animals to obtain the UPR^mt^ per worm. Three biological replicates were performed, with 8 technical replicates per condition.

### 1.11 Mitochondrial membrane potential relative analyses by flow cytometry

N2 Bristol worms were grown from embryo phase on 10 cm NGM plates supplemented with 2 mM IPTG and 100 µg/mL ampicillin seeded with *pdha-1(RNAi)* or L4440 control bacteria until stage L4. Worms were collected with 10 mL S. Basal and transferred onto 6 cm RNAi plates supplemented with 1 µM TMRE and 4 µM MTG for 24 h at 20°C. Worms were subsequently washed twice with 10 mL S. Basal and transferred to RNAi plates for 1h at 20°C to allow intestinal clearing of the fluorescent dyes, and imaged with a CellInsight CX5 high-content imager or analyzed with flow cytometry using a BioSorter (Union Biometrica). For Biosorter analyses, the following excitation wavenegths were used: λ_excitation_ = 488 nm and λ_emission_ = 510 nm. The fluorescence intensity in the red and green channels was normalized by size (time of flight) of the worms (Supplementary Fig. 5). To determine the normalized membrane potential, the fluorescence measured in the red channel was normalized to the fluorescence measured in the green channel.

### 1.12 *C. elegans* extracellular flux analysis

Baseline and maximal oxygen consumption rates (OCR) were assessed using a Seahorse XF Pro Analyzer (Agilent Technologies) in an adaptation of a published protocol ^32^. 24 h prior to the experiment, the XF Pro heater was turned off and the sensors hydrated with 200 µL of XF Calibrant at 37°C in a non-CO_2_ incubator. Micropipette tips were washed with 0.1% Triton-X (v/v) in S. basal to inhibit worms from sticking to the plastic. Wild type (N2 Bristol), *pdha-1(RNAi)* and *dld (RNAi)* knockdown worms, treated with water or 25 mM DCA were collected at L4 stage, were washed 2 times with 1 mL S. basal, diluted with S. basal to 3 worms/10 µL, and transferred in 160 µL NaN_3_ (Sigma Aldrich S2002-25G) dissolved in S. basal at 25 µM and 50 mM final concentrations upon worm exposure, respectively. The optimal concentration of FCCP was determined using a titration curve, using wild type (N2 Bristol), (Supplementary Fig. 4B), *pdha-1(RNAi)* (Supplementary Fig. 4C) and *dld-1(RNAi)* (Supplementary Fig. 11C) worms. The assay timing was 2 min of mixing, 30 s delay, 2 min of measurement, with cycles repeated 4 times for baseline, 7 times for FCCP, and another 4 times for NaN_3_. 15 technical replicates were used per condition for each of three biologically independent experiments run at approximately 21-23°C. Non-mitochondrial O_2_ consumption was not subtracted from baseline and maximal measures, as there was no significant difference between groups and the contribution was minimal (Supplementary Fig. 4E). Statistical analysis was performed in Prism 10.2.2 (GraphPad) on OCR values following normalization to worm count. The average OCR within a measurement period was determined for each well, with median response within a biological replicate taken across 15 technical replicates. Differences between wild type and knockdown worms and between untreated and DCA treated worms were analyzed by Welch’s t-test. Shown are mean +/- standard error of the mean.

### 1.13 Lactate and pyruvate analyte analyses

Wild type (N2 Bristol), *pdha-1(RNAi), dld-1(RNAi), dld-1(1:20 RNAi)* and *dld-1(1:100 RNAi)* worms were grown on 15 cm NGM plates with 0.2 mM IPTG, 100 µg/mL ampicillin, treated with 25 mM DCA or water, from embryo stage until stage L4+1. Animals were then washed with 10 mL S. Basal and collected in 15 mL conical tubes. After washing three times with S. Basal, worms were flash frozen in liquid nitrogen and stored at -80°C. Lactate and pyruvate were spectrophotometrically analyzed as previously described ^33^.

### 1.14 Western immunoblot analyses

Wild type (N2 Bristol), PDHA-1 and DLD-1 knockdown worms were synchronized by bleaching. Stage L4 worms (ten worms per condition) were washed three times with S. Basal buffer and stored on ice. An equal volume of RIPA buffer and 6× Loading Dye (0.3 M Tris-HCl, 12% SDS, 0.06% Bromophenol blue, 50% Glycerol, 0.6 M DTT) was subsequently added, and samples were incubated at 95°C for 15 min. Samples were then centrifuged at 21300*g* for 20 min and the supernatant was loaded on Mini-PROTEAN® TGX™ Precast Protein Gels, 4–15% (Bio-Rad). Proteins were blotted onto polyvinyl difluoride membrane using Wet Transfer Buffer (38.6 mM glycine, 47.9 mM Tris and 20% methanol, pH 8.3) at 4°C for 1 h at 350 mA. Membranes were first blocked in Intercept (TBS) Blocking Buffer (LICORbio) at room temperature for 1 h. The blocked membranes were incubated overnight with primary antibodies at the following dilutions: PDHA1 (Abcam, ab110334, 1:1000 dilution), DLD (Abcam, ab133551, 1:1000 dilution), beta-actin (GeneTex, GTX109639, 1:1000 dilution). Membranes were then washed three times with 1× TBS-Tween20 (TBST; TBS with 0.1% Tween20) for 10 min and incubated with secondary antibodies (IRDye 800CW Anti-Rabbit IgG Goat and IRDye 680CW Anti-Mouse IgG Goat) for 1 h at room temperature. Membranes were then washed at room temperature 3 times for 10 min with TBST, and 2 times for 10 min with TBS, and imaged using an Odyssey CLx infrared imaging system. The relative protein band densities were quantified using ImageJ ^34^. The full gel images for PDHA-1 are in Supplementary Fig. 1B, and the full gel images for DLD-1 are in Supplementary Fig. 7B.

### 1.15 Confocal imaging

Confocal imaging was performed using the *C. elegans* SJ4103 strain carrying the zcls14 construct (*myo-3p::GFP(mit)*) which expresses the GFP in the mitochondria of the worm body-wall muscle. The animals were grown in large numbers and bleached onto RNAi NGM plates prepared as described above, seeded with L4440 (control), *dld-1 (RNAi)*, and *pdha-1 (RNAi)* for 3 or 4 days treated with water or 25 mM DCA. On the day of imaging, the animals were collected and placed on 3% agarose gel pad on a slide in 20-25 μM levamisole dissolved in S-basal. The cover slip was placed on top of the paralyzed worms and sealed to the agarose pad with nail polish. The mitochondria in the body-wall muscle were imaged with excitation at 488 nm and emission range of 493-589 nm at 63× magnification on a confocal microscope to assess mitochondrial morphology.

### 1.16 Statistical analyses

Statistical analyses were performed using GraphPad Prism version 10.2.2. Unless otherwise noted, differences between wild type and knockdown worms were analyzed by Welch’s t-test while comparison of untreated knockdown worms and the dose response to DCA were analyzed by one-way ANOVA followed by post-hoc analysis for a log linear regression trend if the ANOVA p value was significant. Statistical significance is reported for the ANOVA along with the estimated slope ± standard error (SE) of the slope. Data for individual conditions are presented by mean ± standard deviation (SD). For studies of survival, Kaplan Meier survival curves were plotted and compared with a log-rank Mantel Cox test. Statistical significance was set at p value < 0.05 with a Bonferroni correction applied to account for multiple testing when needed. For the automated light stimulated locomotor activity, comparisons between groups were analyzed using ANOVA with a post-hoc Tukey test. An adjusted probability (P) value of less than 0.05 was considered statistically significant. Plots were made using ggplot2 ^29^.

## 2 Data availability

## 3 RESULTS

### DCA treatment significantly rescued disease phenotypes in pdha-1(RNAi) worms

The most common cause (∼77%) of PDCD is pathogenic variants in the PDHc alpha subunit gene (*PDHA1*)^9^. Therefore, we first characterized the effect of knocking down the ortholog, *pdha-1,* in *C. elegans* by feeding wild-type (N2 Bristol) worms with RNAi bacteria. We confirmed a 70% reduction of PDHA-1 protein in the *pdha-1(RNAi)* knockdown worms using western immunoblotting (Fig. 1A, 1B, Supplementary Fig 1B). Having established a *C. elegans* model for PDHA-1 deficiency, similar to that developed previously^23^, we investigated the effect of human clinical trial grade DCA on their disease phenotypes. Wild-type and *pdha-1(RNAi)* worms were treated from embryo phase on solid media supplemented with a 5-point log concentration dosing range of DCA (10 µM, 100 µM, 1 mM, 5 mM, 25 mM); animals were subsequently observed at L4+1 and L4+5 days of adulthood to visually identify toxicity at the level of gross morphologic alterations and determine the maximal safe effective dosage exposure (Supplementary Fig. 2, Supplementary Fig. 3). No toxic effects on gross morphology were observed in the tested DCA dosage range.

**Fig. 1.**
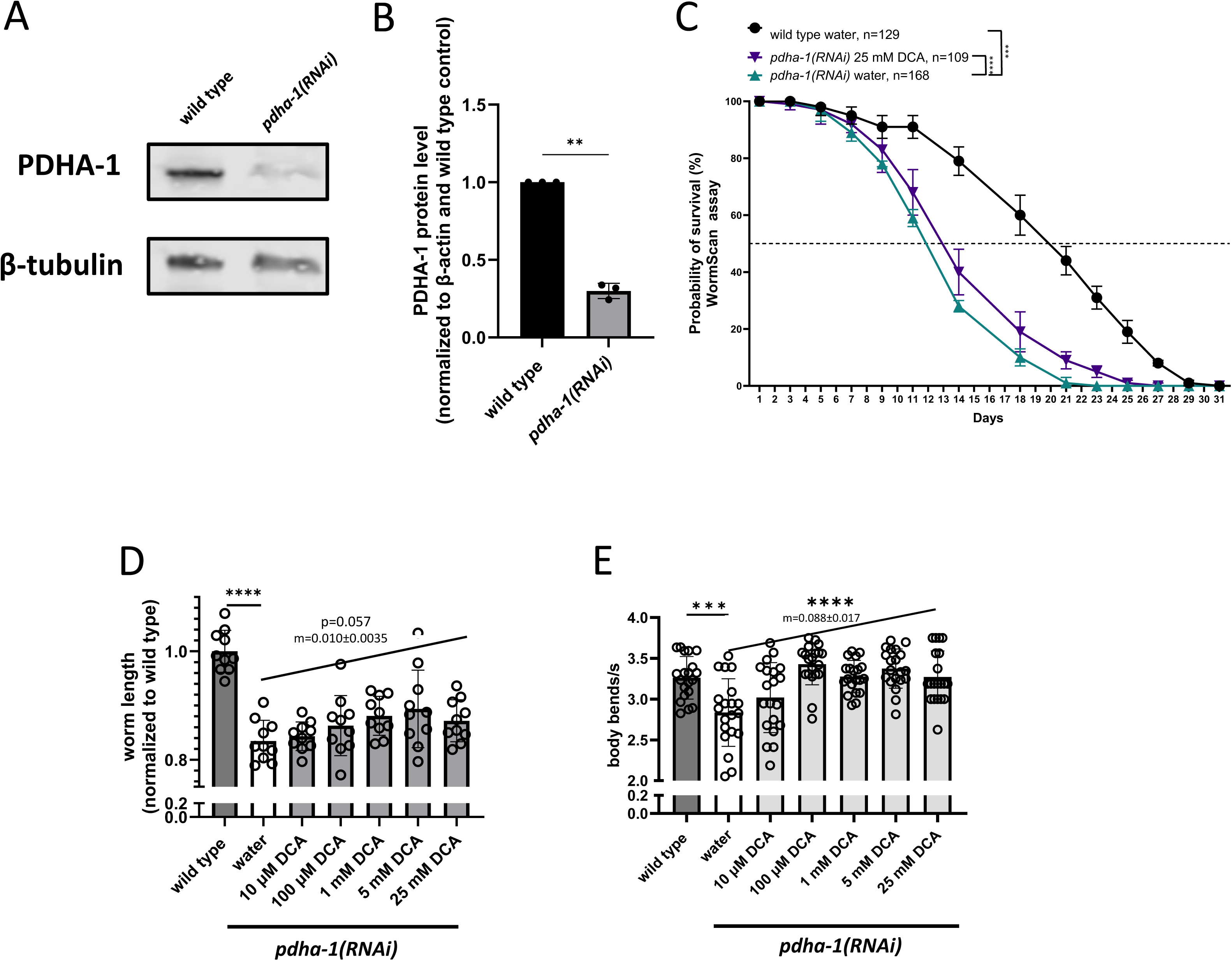
DCA treatment effect on overall animal health of *pdha-1(RNAi)* knockdown worms. **(A)** Western blot analysis of PDHA-1 protein levels after feeding N2 (Bristol) worms with *pdha-1* or empty vector *RNAi* bacteria. A representative blot of n=3 biological replicates is shown. **(B)** Quantification of the PDHA-1 protein bands density, confirming the specific knockdown of PDHA-1 by feeding worms with *pdha-1(RNAi).* PDHA-1 protein levels relative to β-actin were normalized to wild type. Bars represent the mean ± SD of 3 biological replicates. **(C)** Lifespan analysis of the *pdha-1(RNAi)* knockdown worms (teal curve) in the presence of FUDR shows a significant decrease in lifespan relative to wild type (N2 Bristol, black curve, p = 0.0083), while treatment with 25 mM DCA significantly increased worm survival (purple curve, p<0.0001). A representative trial of 3 biological replicates is presented. **(D)** Worm growth analysis of the *pdha-1(RNAi)* knockdown worms (white bars) show a ∼12% decrease in length compared to wild-type worms (N2 Bristol, dark grey bars, **** p<0.0001), while treatment with 10 µM to 25 mM DCA (light grey bars) leads to a strong dose-dependent increasing trend in worm length (one-way ANOVA, p=0.057, post hoc log linear trend shown). Data represent the mean ± SD of 10 individual animals. **(E)** *pdha-1(RNAi)* knockdown worms (white bars) had reduced swimming (neuromuscular) activity relative to wild- type control worms (N2 Bristol, dark grey bar, *** p < 0.001), which was significantly rescued by treatment with 10 µM to 25 mM DCA (light grey bars, one-way ANOVA, **** p < 0.0001, log linear trend shown). Treatment with 100 µM, 1 mM, 5 mM and 25 mM DCA lead to a recovery not significantly different than wild-type swimming activity. Data represent the mean value ± SD of 20 individual animals.

Lifespan of the *pdha-1(RNAi)* worms was first quantified by our previously established WormScan method performed in the presence of FUDR ^35^, where we observed a significant reduction in survival (p = 0.0083) relative to wild-type control (N2 Bristol) worms (Fig. 1C). Treatment with 25 mM DCA treatment significantly increased lifespan of the *pdha-1(RNAi)* worms, with the median lifetime increased from 12 to 15 days, p < 0.0001 (Fig. 1C). No effect was observed on lifespan of the wild-type worms after treatment with 25 mM DCA (Supplementary Fig. 1C). Furthermore, the effect of DCA on *pdha-1(RNAi)* worms was independently confirmed using a standard manual lifespan assay without FUDR (Supplementary Fig. 1D), in which treatment with 25 mM DCA significantly increased lifespan compared to untreated *pdha-1(RNAi)* worms (p= 0.013). In addition, worm growth was quantified in *pdha-1(RNAi)* knockdown worms, which had an ∼18% decrease in worm length compared to wild-type animals (Fig. 1D). The short growth phenotype showed a trend of rescue by DCA; one-way ANOVA p=0.057, with a linear slope of 0.010 ± 0.003 (Fig. 1D). Post-hoc analysis showed a significant difference in linear length between untreated *pdha-1 (RNAi)* animals and those treated with 5 mM DCA (p=0.0369). The linear trend detection of the ANOVA analysis indicated the slope was ∼3× its standard error (0.010 to 0.0035), which has been further confirmed by log-linear regression of normalized worm length to log[DCA] where the slope is 0.012 ± 0.004 (Supplementary Fig. 1F).

Since PDHc deficiencies cause neuromuscular disease in human patients ^2^, we analyzed the neuromotor activity of the *pdha-1(RNAi)* worms at day 1 adult stage using a swim-locomotor (animal thrashing) assay of worms placed in liquid media ^30^. Indeed, knocking down PDHA-1 significantly decreased worm activity, p< 0.001. Treatments with DCA from 100 µM to 25 mM each significantly improved *pdha-1(RNAi)* worm swim activity by one-way ANOVA, p<0.0001, with a linear response slope 0.088±0.017 (Fig. 1E). Importantly, an activity level not significantly different from wild type was observed with all treatments above 100 µM DCA. In addition, we measured worm light-stimulated activity on solid media, as previously described ^28^. *pdha-1(RNAi)* worms exhibited reduced activity compared to the wild-type control, which was not altered by treatment with DCA at any tested concentration across six technical replicates (Supplementary Fig. 1E). In contrast, DCA treatment increased locomotor activity in wild-type animals (Supplementary Fig. 1E).

We also sought to determine if treatment with DCA improved mitochondrial health at the level of integrated UPR^mt^ induction in *pdha-1(RNAi)* worms. UPR^mt^ has been extensively characterized in *C. elegans*, where it triggers the upregulation of the mitochondrial chaperones *hsp-6* (orthologue to human HSP70) and *hsp-60* (orthologue to human HSP60, or HSPD1) to promote protein folding and maintain mitochondrial health. UPR^mt^ stress has been shown to be triggered by perturbations in mitochondrial physiology ^36,37^ ^38^. Therefore, activation of the *hsp-6p* reporter was used as a quantitative indicator of mitochondrial dysfunction ^39,40^. For these assays, we used an N2 Bristol strain carrying an *hsp-6p::gfp* reporter (green fluorescence) stably crossed to a *myo-2p::mCherry* reporter line (red fluorescence), to quantify the degree of UPR^mt^ stress induction normalized per worm (Supplementary Fig. 4A). The *pdha-1(RNAi)* worms had significantly increased mitochondrial stress induction relative to wild-type animals, which DCA treatment significantly reduced in all tested dose conditions (Fig. 2A). The greatest degree of UPR^mt^ stress reduction (86%, p=0.0012) was observed after treatment with 25 mM DCA.

**Fig 2.**
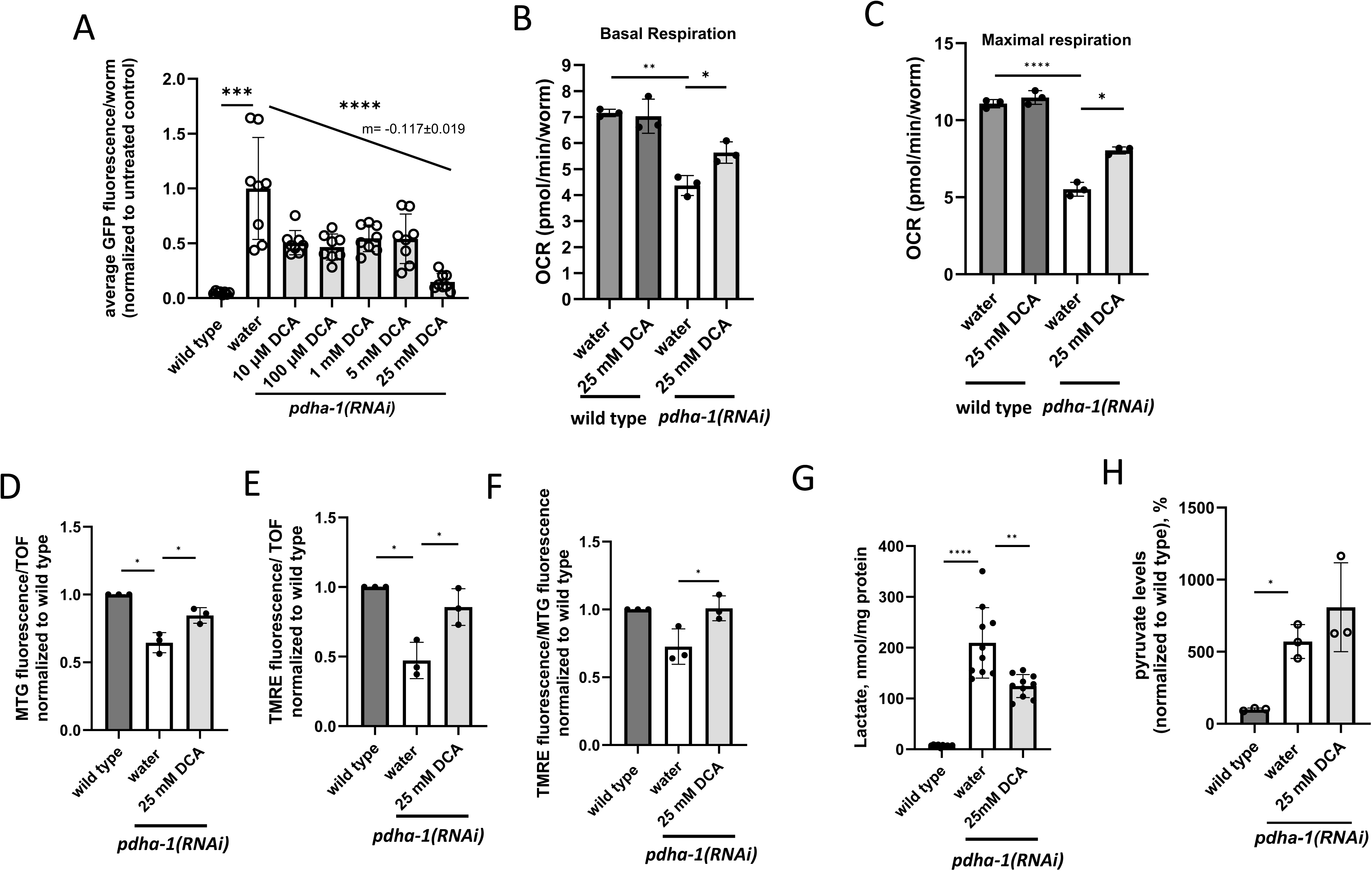
DCA treatment effect on mitochondrial physiology of *pdha-1(RNAi)* knockdown worms. **(A)** Untreated *pdha-1(RNAi)* knockdown worms (white bars) show a significant increase in mitochondrial stress relative to the empty vector control (*hsp-6p::GFP; myo-2p::mCherry*, dark grey bar, *** p < 0.001). Treatment with a 4-log range of DCA concentrations protected animals from UPR^mt^ stress induction (light grey bars, 1-way ANOVA p < 0.0001, post-hoc linear trend indicated). Data represent mean ± SD of 8 replicates with ∼50 animals per replicate and condition. **(B, C)** Basal (B) and maximal (C) mitochondrial respiratory capacity measured by Seahorse assay were significantly reduced in buffer-treated *pdha-1(RNAi)* worms relative to wild type (N2 Bristol). Treatment with 25 mM DCA significantly increased mitochondrial respiration rates in *pdha-1(RNAi)* knockdown animals, while no significant effect was observed in wild-type controls. Data are presented as mean ± SD from three biological replicate experiments. **(D)** Mitochondrial content, measured as relative MTG fluorescence normalized to time of flight, was significantly decreased in untreated *pdha-1(RNAi)* knockdown worms compared to wild- type control (N2, Bristol) and significantly increased following DCA treatment. Data represent mean ± SD of three biological replicate experiments. **(E)** Mitochondrial membrane potential, measured as relative TMRE fluorescence normalized to time of flight, was reduced in untreated *pdha-1(RNAi)* knockdown worms compared to wild-type (N2 Bristol) and was restored upon treatment with 25 mM DCA. Data represent mean ± SD of three biological replicates per condition. **(F)** Relative membrane potential, defined as TMRE fluorescence divided by MTG fluorescence, was decreased in *pdha-1(RNAi)* knockdown worms relative to wild-type (N2 Bristol) and increased after treatment with 25 mM DCA. Data convey mean ± SD of three biological replicate experiments per condition. **(G)** Tissue lactate levels were significantly increased (43%) in the untreated *pdha-1(RNAi)* knockdown worms relative to wild-type (N2 Bristol) and significantly decreased after treatment with 25 mM DCA (p=0.0037). Data represent mean ± SD of 10 biological replicates per condition of 1,000 animals each. **(H)** Tissue pyruvate levels were significantly increased in *pdha-1(RNAi)* knockdown worms compared to wild type (N2 Bristol) both with and without treatment with 25 mM DCA. Bars represent mean ± SD of 3 biological replicates per condition of 1,000 animals each. *: p <0.05, **: p<0.01, ****: p<0.0001.

To further assess the impact of DCA on mitochondrial physiology, Seahorse-based respiration assays were performed on *pdha-1(RNAi)* and wild-type worms treated from embryo stage with either water or 25 mM DCA. Consistent restoration of both basal (Fig. 2B) and maximal (Fig. 2C) respiration was seen in DCA-treated *pdha-1(RNAi)* worms across three biological replicate experiments. No significant changes in mitochondrial respiration were detected after treatment of wild-type worms with 25 mM DCA (Fig. 2B, C). Individual OCR traces for all conditions are provided in Supplementary Fig. 4D. Additionally, the effect of DCA treatment was studied on mitochondrial content and relative mitochondrial membrane potential in the *pdha-1(RNAi)* worms using BioSorter flow cytometry. Specifically, animals treated with either water (buffer control) or 25 mM DCA were co-stained with MitoTracker Green (MTG) and tetramethylrhodamine ethyl ester (TMRE) to quantify mitochondrial content and membrane potential, respectively. Knockdown of PDHA-1 led to a significant mean reduction in both mitochondrial content (36%, Fig. 2D) and mitochondrial membrane potential (53%, Fig. 2E) as compared to wild-type worms. Importantly, treatment with 25 mM DCA significantly restored both mitochondrial mean parameters to near wild-type levels (Fig. 2B, 2C, Supplementary Fig. 5). Similarly the relative mitochondrial membrane potential normalized for mitochondrial content, calculated based on TMRE/MTG fluorescence ratio, showed a trend toward reduction in *pdha-1(RNAi)* worms, which significantly increased following DCA treatment (Fig. 2F). Validation of these alterations in mitochondrial physiology was performed by an independent method involving quantitation of MTG and TMRE relative fluorescence in these animals using high-content microscopy (CellInsight CX5). This imaging-based analysis revealed the same overall trends as were observed by flow cytometry, further supporting the conclusion that both mitochondrial content and mitochondrial membrane potential are reduced in *pdha-1(RNAi)* worms and restored by DCA treatment (Supplementary Fig. 6A–D). In addition, we assessed mitochondrial morphology by confocal microscopy. *pdha-1 (RNAi)* animals showed abnormal and disorganized mitochondrial networks compared with wild-type animals (Supplementary Fig. 6F). Treatment with 25 mM DCA improved overall network organization, with mitochondria appearing more evenly distributed and forming more continuous filamentous structures with reduced clustering (Supplementary Fig. 6F). PDHc catalyzes the conversion of pyruvate to acetyl-coenzyme A, which enters into the tricarboxylic acid (TCA) cycle. Reduced PDHc activity leads to pyruvate accumulation, which is subsequently reduced to lactate through the lactate dehydrogenase (LDH) equilibrium reaction. Accordingly, PDCD patients commonly exhibit elevated plasma lactate and pyruvate levels. We therefore measured these metabolites in tissues from *pdha-1(RNAi)* knockdown worms to determine whether similar metabolic alterations occur. As anticipated, lactate (Fig. 2G) and pyruvate (Fig. 2H) metabolite levels were significantly increased in the *pdha-1(RNAi)* whole worm populations. Interestingly, 25 mM DCA treatment throughout worm development caused a significant decrease in tissue homogenate lactate in *pdha-1(RNAi)* worms (p=0.0037, Fig. 2G), but did not significantly alter the *pdha-1(RNAi)* tissue homogenate level of pyruvate (Fig. 2H). While the pyruvate:lactate ratio was increased in *pdha-1(RNAi)* worms p<0.05, 25 mM DCA treatment did not significantly rescue this biochemical ratio (Supplementary Fig. 6E), as is consistent with no anticipated effect of DCA on the NAD^+^/NADH cytosolic redox ratio.

### 3.1 DCA treatment significantly rescued growth, neuromuscular activity and survival in dld-1(RNAi) worms

DLD-1 encodes the E3 subunit of several keto acid dehydrogenases, including PDHc that has been previously modeled by RNAi knockdown in *C. elegans* ^11,41^ with varying phenotypic severity directly correlating with the degree of RNAi-induced DLD-1 knockdown. To model the therapeutic efficacy of DCA on a wide range of severity of PDHc deficiency in *C. elegans*, a graded dilution of *dld-1(RNAi)* bacteria with bacteria harboring an L1440 control vector were used at ratios of 1:100 and 1:20. The varying degree of DLD-1 knockdown by RNAi dilution was confirmed by the correlated reduction of DLD-1 protein expression in whole worm lysates by western immunoblotting (Fig. 3A, B, Supplementary Fig. ***7***B). As described above, treatment with DCA was performed from embryo phase, and worms were monitored for potential toxicity at stage L4+1 day and L4+5 days of adulthood. No toxic effects were observed (Supplementary Fig. 8, Supplementary Fig. 9). Similar to previous reports ^11,41^, lifespan analyses showed that *dld-1(RNAi)* knockdown worms had increased lifespan, as measured in N2 Bristol by the WormScan method in the presence of FUDR (Fig. 3C) and a standard manual lifespan assay without FUDR (Supplementary Fig. 10A). Additionally, the *dld-1(1:20 RNAi)* knockdown strain showed a significant decrease in lifespan relative to the wild-type control worms as measured by both WormScan (Fig. 3D, p<0.0001) and manual lifespan assay (Supplementary Fig. 10A, p=0.002). In contrast, the *dld-1(1:100 RNAi)* worms showed significantly decreased lifespan when assessed using the WormScan method (Supplementary Fig. 10B, p<0.001), whereas no significant change in lifespan relative wild type control was observed using the standard manual lifespan assay (Supplementary Fig. 10C). Interestingly, 25 mM DCA treatment trended to increase survival in *dld-1(RNAi)* (Fig. 3C, Supplementary Fig. 10D) and significantly extended survival in the *dld-1(1:20 RNAi)* worms (Fig. 3D, Supplementary Fig. 10A), but did not alter lifespans of the *dld-1(1:100 RNAi)* strain (Supplementary Fig. 10B, C). Thus, a survival benefit from DCA treatment was seen in animals with a more severe inhibition of DLD expression, consistent with our previou***s*** report using laboratory-grade DCA^11^.

**Fig. 3.**
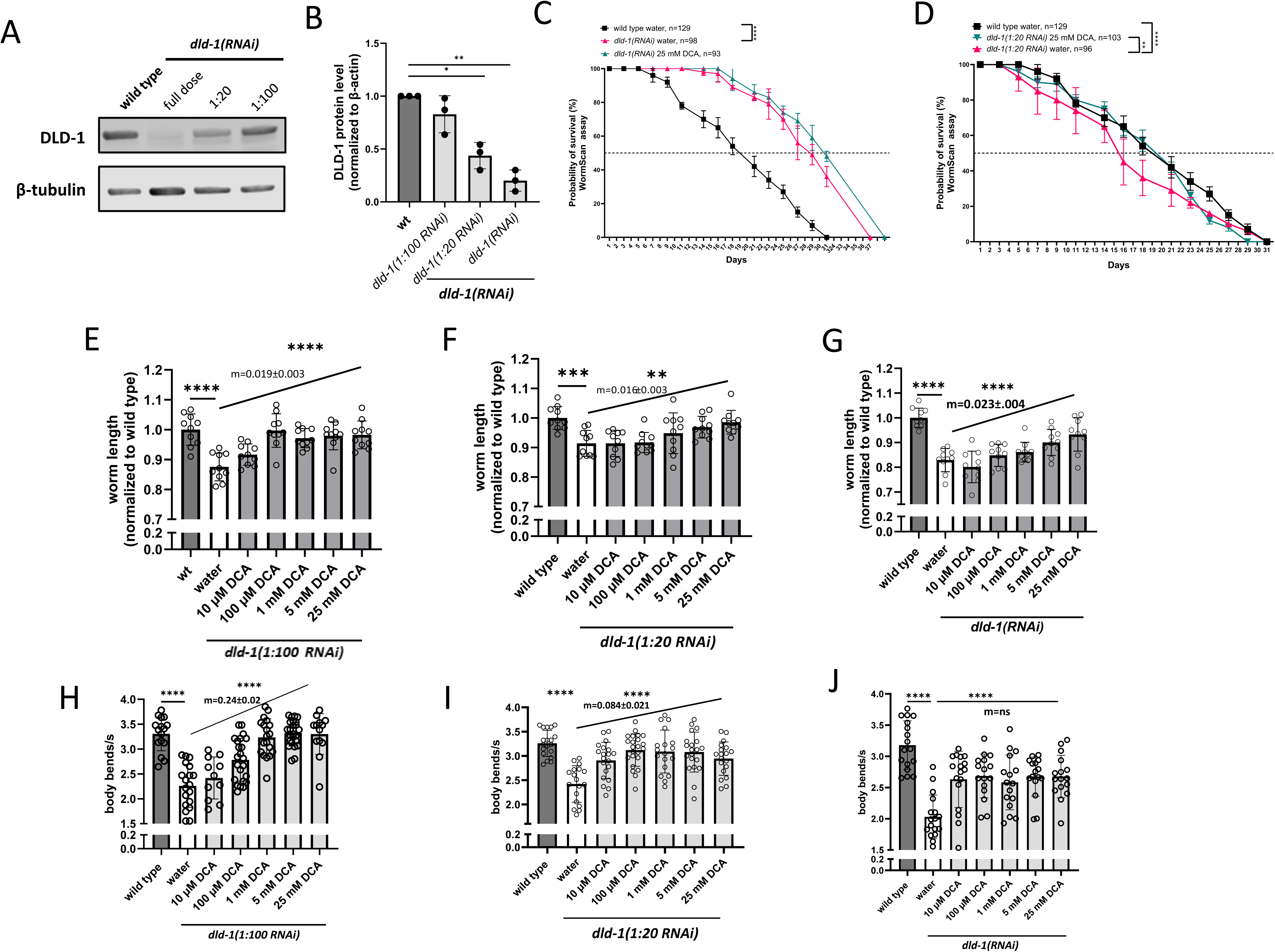
DCA treatment effect on overall animal health of *dld-1(RNAi)* graded severity knockdown worms. **(A)** Western blot analysis of DLD-1 protein levels in *C. elegans* fed with *dld-1(RNAi)* bacteria (undiluted, 1:20 and 1:100 dilution with L1440) and L1440 (wild-type, N2 Bristol). A representative blot of n=3 biological replicates is presented. (**B)** Quantification analysis of DLD-1 protein band densities in DLD-1 knockdown worms shows that feeding worms with increasing concentrations of *dld-1(RNAi)* bacteria leads to a consistent decrease in DLD-1 protein levels in total worm lysates relative to wild type (N2 Bristol). The levels of DLD-1 relative to β-actin were normalized to the wild-type control. Data represent the mean ± SD of 3 biological replicates. **(C)** Lifespan analysis of the *dld-1(RNAi)* worms in the presence of FUDR shows a significant increase (pink curve, p<0.0001) relative to the empty vector control (N2 Bristol, black curve). Treatment with 25 mM DCA caused a trend toward increased lifespan (teal curve). A representative trial of 3 biological replicates is presented. **(D)** Lifespan analysis of the *dld-1 (1:20) RNAi* worms (pink curve) in the presence of FUDR shows a significant decrease in survival relative to the wild type worms (N2 Bristol, black curve, p=0.018), whereas treatment with 25 mM DCA caused a significant increase in lifespan (teal curve, p<0.0001). A representative trial of 3 biological replicates is presented. **(E-G)** Worm growth analysis of *dld-1(RNAi)* knockdown worms showed a significant decrease in worm length for all *dld-1(RNAi)* bacteria dilutions tested (white bars) relative to wild type (N2 Bristol). Treatment with DCA significantly rescued growth in all *dld-1(RNAi)* knockdown worms. All three RNAi dilutions showed a significant dose-dependent log linear response by 1-way ANOVA. Bars represent the mean ± SD of 10 individual animals, the slope of the linear regression ± Std Er is shown. Post-hoc multiple comparisons showed for *dld-1(1:100 RNAi)* that every DCA dose except 10 µM demonstrated rescue p<0.0001 (E). For *dld-1(1:***2***0 RNAi)* the 5 mM and 25 mM DCA doses demonstrated significant rescue, p=0.042 and p=0.0048, respectively (F) and *dld-1(RNAi)* the 5 mM and 25 mM DCA doses demonstrated significant rescue, p=0.019 and p=0.0003, respectively (G). **(H-J)** *dld-1(RNAi)* knockdown worms had significantly decreased worm swimming (neuromuscular) activity (white bars) relative to wild-type (N2 Bristol, dark grey bars), while treatment with DCA significantly improved this phenotype (light grey bars) at all doses tested. *dld-1*(*1:100 RNAi*) and *dld-1*(*1:20 RNAi*) knockdown worms showed a significant dose-dependent linear response, with a lower slope for the 1:20 RNAi dilution. With the undiluted RNAi dose, significant rescue was observed but no dose dependence was detected. Bars represent the mean value ± SD of 20 individual animals. *: p <0.05, **: p<0.01, ***: p<0.001, ****: p<0.0001.

*dld-1(RNAi)* knockdown caused a consistent linear growth defect in all three *dld-1* knockdown concentrations tested (Fig. 3E-3G). DCA treatment demonstrated significant dose-dependent rescue of this phenotype by one-way ANOVA analysis of *dld-1(1:100 RNAi), dld-1(1:20 RNAi)* and *dld-1(RNAi)*, p <0.0001, p<0.01 and p<0.0001, respectively, (Fig. 3E-3G). A two-way ANOVA analysis of these data indicated a highly significant (p < 0.0001) effect on worm length occurs based both on DCA dose and RNAi dose. *dld-1(RNAi)* knockdown impact on worm neuromuscular activity was monitored by thrashing analysis in liquid culture, where their body bends/s were quantified. Significantly decreased (p<0.0001) thrashing activity by ∼35% was observed in all *dld-1(RNAi)* knockdown strains relative to wild-type worms (Fig. 3H-3J). DCA treatment significantly improved thrashing activity in all three *dld-1(RNAi)* knockdown strains. One-way ANOVA analysis of DCA treatment with post-hoc analysis of a log linear regression trend showed that the dependence on DCA concentration directly correlated with degree of dilution, where DCA treatment had a greater beneficial effect on rescuing worm neuromuscular activity at the mildest (1:100 dilution) than moderate (1:20 dilution) degree of DLD-1 inhibition, and no significant dose dependence was seen at the most severe degree of full-dose *dld-1(RNAi)*. Similar to the worm length analysis presented above, a two-way ANOVA of the neuromuscular activity data revealed a strong dependence (p < 0.0001) on both RNAi concentration and DCA treatment concentration. In addition, we used an automated assay to assess worm light stimulated activity at days 2, 5 and 10 of adulthood, as described above. Untreated *dld-1(full-dose RNAi)* worms exhibited an abnormal increased activity compared to wild-type controls, while treatment with 100 µM or 1 mM DCA reduced locomotor activity toward wild-type levels (Supplementary Fig. 10E).

### DCA treatment effect on mitochondrial stress induction correlated with the severity of the disease phenotype in the graded RNAi dilution of dld-1(RNAi) knockdown worms

Similarly, as described for *pdha-1(RNAi)*, mitochondrial stress induction was studied in *dld-1(RNAi)* knockdown strains in a background of N2 Bristol worms stably carrying two transgenic reporters: *hsp-6p::gfp* (green) and *myo-2p::mCherry* (red) (Supplementary Fig 11A). These double-transgenic worms were fed three different graded RNAi dilutions, *dld-1(1:100 RNAi), dld-1(1:20 RNAi)* and *dld-1(RNAi)* bacteria from embryo phase, with UPR^mt^ stress induction analyzed at stage L4+1 Day (first day of adulthood) and normalized to the number of worms detected by mCherry fluorescence. As expected, we observed a dramatic increase in UPR^mt^ stress at all three dilutions of *dld-1(RNAi)* bacteria tested (Fig. 4A-4C) p<0.0001. The greatest degree of *hsp-6p* dependent induction was seen in full-dose RNAi knockdown, with proportionate decreased induction in moderate (1:20) and mild (1:100) RNAi induction (Supplementary Fig. 11B). One-way ANOVA analysis of the DCA effect with post-hoc analysis for a log linear trend revealed that DCA did not alter the UPR^mt^ stress induction level with full dose *dld-1(RNAi)* knockdown (Fig 4C), but a significant dose-response rescue was observed for the moderate *dld-1(1:20 RNAi),* p<0.001, and mildest *dld-1(1:100 RNAi),* p<0.0001, knockdown strains (Fig 4BA, respectively), with the DCA dose response being greater for *dld-1(1:100 RNAi)*. Representative images showing the effect of DCA treatment on mitochondrial unfolded protein response stress are presented in Supplementary Fig. 11A. As described above, mitochondrial respiration was evaluated in *dld-1(full-dose RNAi)* animals treated with either water or 25 mM DCA. The *dld-1(RNAi)* knockdown worms displayed a significant reduction in both basal and maximal respiration relative to the wild-type control (Fig. 4D, E). Treatment with 25 mM DCA significantly improved both basal and maximal respiratory capacity, suggesting a partial restoration of mitochondrial function (Fig. 4D, E). Full mitochondrial respiration plots for each condition are shown in Supplementary Fig. 11D. Lactate levels were incrementally greater in full-dose *dld-1(RNAi)* relative to wild-type worms compared to either dilution strain. Confocal imaging *dld-1* (RNAi) worms revealed the appearance of short branches extending from irregular mitochondrial structures with reduced network connectivity relative to wild type. Upon treatment with 25 mM DCA, mitochondrial morphology showed increased continuity and filament formation, with fewer isolated puncta and a more connected network (Supplementary Fig. 6F).

**Fig. 4.**
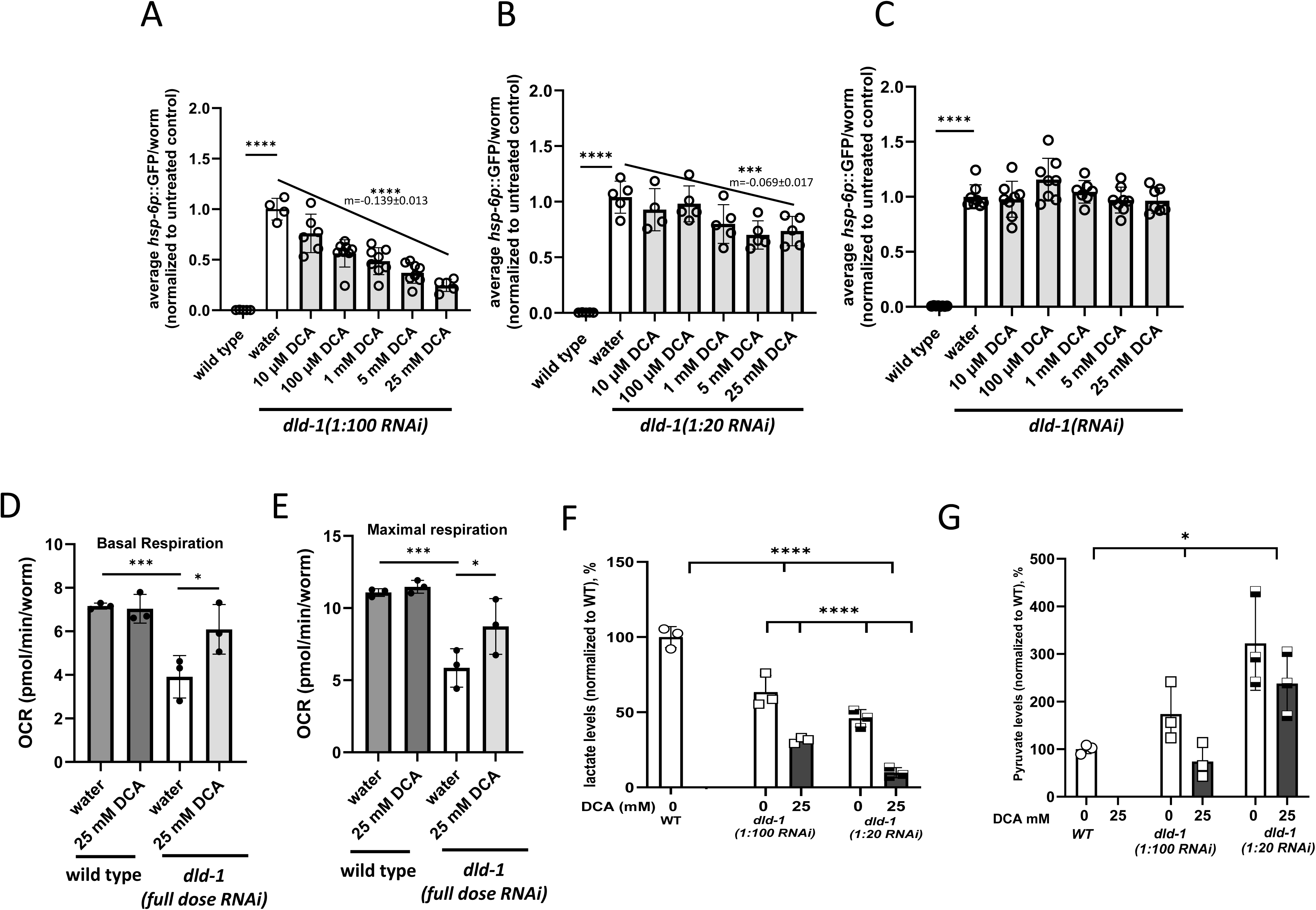
DCA treatment effect on mitochondrial physiology of *dld-1(RNAi)* graded severity knockdown worms. **(A-C)** All *dld-1(RNAi)* knockdown worms (white bars) showed significantly greater induction of the mitochondrial stress response compared to wild-type control (*hsp-6p::GFP; myo-2p::mCherry*, dark grey bars; ****, p<0.0001). Treatment with DCA significantly reduced mitochondrial stress for *dld-1*(*1:100 RNAi*) and *dld-1*(*1:20 RNAi*) by one-way ANOVA, with the log linear regression slope increased from -0.139 to -0.069. With the undiluted *dld-1*(*RNAi*) no significant effect on mitochondrial stress was observed. Bars represent the mean ± SD of 8 replicates with ∼50 animals each. **(D, E)** Oxygen consumption rate (OCR) measured by Seahorse assay reveals reduced basal **(D)** and maximal **(E)** respiration in untreated *dld-1*(full-dose RNAi) worms relative to wild-type (N2 Bristol), whereas treatment with 25 mM DCA significantly increased respiration rates. No effect was observed following treatment of wild-type worms with 25 mM DCA. Data are presented as mean ± SD from three biological replicate experiments. **(F)** Tissue lactate levels were significantly decreased in *dld-1(1:20 RNAi)* and *dld-1(1:100 RNAi)* knockdown worms (white bars) compared to the empty vector control (N2 Bristol, dark grey bars). Further reduction in lactate levels occurred after treatment with 25 mM DCA. Data represent mean ± SD of 3 biological replicates of 1,000 animals each. **(G)** Tissue pyruvate levels in the *dld-1* knockdown worms (white bars) trended toward increased in *dld-1(1:20 RNAi)* by 322% (p=0.058) and in *dld-1(1:100 RNAi)* by 174% (p=0.063) relative to wild-type levels (N2 Bristol, dark grey bars). Treatment with 25 mM DCA significantly decrease pyruvate tissue levels (black bars). *: p <0.05, **: p<0.01, ***: p<0.001, ****: p<0.0001.

Surprisingly, lactate levels were significantly greater in the mildest (1:100) than moderate (1:20) RNAi dilution worms. 25 mM DCA significantly reduced lactate levels in 1:100 and 1:20 dilution strains of *dld-1(RNAi)* knockdown (Fig. 4F). Pyruvate levels showed an opposite trend, with greater pyruvate levels relative to wild-type worms evident in a stepwise decreasing fashion in moderate *dld-1(1:20 RNAi),* than mild *dld-1(1:100 RNAi)*, and lowest in full-dose *dld-1(RNAi)* (Fig. 4G). Similarly, pyruvate to lactate ratio was only increased in *dld-1(1:20 RNAi)* worms but not in the *dld-1(1:100 RNAi)* worms (Supplementary Fig. 11E). 25 mM DCA showed non-significant trends toward reduced pyruvate levels in the *dld-1(1:100RNAi)* strain (p=0.085) (Fig 4F) and increased pyruvate:lactate ratio in the moderate *dld-1(1:20 RNAi)* worms (Supplementary Fig 11E). As full-dose *dld-1(RNAi)* did not have increased mitochondrial stress with DCA treatment, lactate and pyruvate levels were not measured in these conditions.

## 4 DISCUSSION

PDHc deficiency (PDCD) is an inherited mitochondrial disorder involving lactic acidosis, neurodevelopmental disability, and high mortality, with treatments largely limited to symptomatic management ^42^ and clinical trial investigation of DCA ^22^. To objectively evaluate the preclinical efficacy of DCA treatment in PDCD, we created and characterized *C. elegans* models of two different genetic causes of PDHc deficiency by feeding RNAi to knock-down expression of *PDHA-1* and *DLD-*1 homologues. Further, *DLD-1* deficiency was modeled with three different severity grades by titrating the concentration of *dld-1(RNAi)* bacteria: full dose (severe), 1:20 (moderate) and 1:100 (mild) dilutions. These worm models recapitulate key phenotypic features observed in PDHc patients, including reduced survival, delayed growth, locomotor impairments, and intermediary metabolic abnormalities including elevated lactate and/or pyruvate tissue levels. Extensive phenotypic analyses of both RNAi knockdown models’ impact on lifespan, health span, intermediary metabolism, and diverse read-outs of mitochondrial physiology were performed at baseline relative to wild-type worms, and following treatment with DCA across a 5-point log-scale (10 µM to 25 mM) in growth, neuromuscular function, and mitochondrial stress induction studies and high-dose (25 mM) DCA in tissue metabolite and lifespan studies. Across all studies, DCA showed no toxic effects at the level of survival, health span (growth, neuromuscular activity), or metabolism (mitochondrial UPR^mt^ stress induction, mitochondrial amount or membrane potential, mitochondrial respiratory capacity, or lactate and pyruvate levels). Indeed, DCA treatment led to statistically significant therapeutic benefit on multiple phenotypic and metabolic outcome measures in both PDHc knockdown *C. elegans* models, including mitochondrial content, membrane potential, respiratory capacity (OCR), stress induction, and morphology, with multiple DCA dose-response effects seen.

Decreasing *pdha-1* expression in *C. elegans* significantly reduced survival, significantly decreased worm linear growth and neuromuscular activity as well as significantly increased mitochondrial stress induction as well as tissue levels of lactate and pyruvate. Additionally, the *pdha-1* knockdown worms showed decreased respiration rates, mitochondrial content and membrane potential. DCA treatment showed statistically significant benefit on animal survival, overall animal health at the level of neuromuscular function (thrashing) and linear growth. In addition, improvements in overall metabolic and mitochondrial physiology upon DCA treatment were evident in the *pdha-1(RNAi)* strain, including reduced lactate, reduced mitochondrial stress induction, increased mitochondrial membrane potential and mitochondrial content, and enhanced basal and maximal mitochondrial respiratory capacity. Interestingly, while DCA effect leveled off in some experimental outcomes, no specific dose threshold was universally beneficial in all conditions studied. Survival analyses were performed following treatment with the highest tested DCA dose (25 mM), since this concentration consistently produced beneficial effects across all other assays, including worm length, neuromuscular activity, and notably, reduced UPR^mt^ stress levels to those observed in wild-type worms. Previously, our research group had studied the intermediary metabolism effects of PDHc RNAi knockdown using stable isotope and mass spectrometry studies^23^, further adding to the body of informative data confirming the relevance of this *C. elegans pdha-1(RNAi)* knockdown model to study PDHA1- based human disease. Treatment with 25 mM DCA in graded models of *dld-1 RNAi* silencing significantly prolonged animal survival only in the more severe knockdown conditions, namely *dld-1(RNAi) and dld-1 (1:20 RNAi)*, similarly as previously reported ^11,41^, with no effect seen in the *dld-1(1:100 RNAi)* dilution strain. Direct comparison of the lifespan results obtained in the present study with those reported in our previous study on PDHc deficiency models^11^ was presented in Supplementary Table 1. Interestingly, our prior work has found that in multiple mitochondrial disease models including in *dld-1(RNAi)* worms, therapies started during development induce a greater degree of outcome rescue than are those started in adult-stage animals. *dld-1(RNAi)* knockdown significantly reduced worm linear growth and neuromuscular activity in all three knockdown *dilutions* tested, with increased induction of mitochondrial stress that inversely correlated with the level of residual DLD-1 protein. However, *dld-1(1:20 RNAi)* and *dld-1(1:100 RNAi)* strains showed a trend towards decreased tissue lactate levels and increased pyruvate. Analysis of DCA treatment in the three *dld-1(RNAi)* knockdown strains confirms a differential sensitivity to DCA that correlated to the severity of the observed phenotype. While treatment with DCA rescued animal growth and neuromuscular activity in all *dld-1(RNAi)* strains, we observed distinct responses of UPR^mt^ stress that correlated to the degree of *dld-1(RNAi)* knockdown. The most severe full-dose *dld-1(RNAi)* knockdown strain showed no improvement with DCA, whereas the less severe *1:20* and *1:100 dld-1(RNAi)* knockdown strains showed an increasing dose-dependent response after treatment with DCA at tested concentrations. However, treatment with 25 mM DCA restored both basal and maximal mitochondrial respiratory capacity in the *dld-1 (RNAi)* worms. Additionally, the *dld-1(RNAi)* animals displayed altered mitochondrial network morphology, as indicated by an increased branch length. Treatment with 25 mM DCA significantly reduced branch length, suggesting a partial improvement in mitochondrial organization. These data add to the preclinical literature in PDHc models of DCA, a structural analog of pyruvate that has been previously shown to rescue disease phenotypes in some PDHc models ^11^ while having no effect in others ^15,21^ ^12^. The reason for this differential effect was proposed to be the nature of the E1 alpha subunit mutations, with mutations affecting the stability of the E1 alpha subunit being responsive, while mutations that affect the catalytic activity of the protein being unresponsive to DCA treatment ^15^. Our prior studies in a very severe zebrafish DLD knockout model that had very low residual PDHc activity and larval mortality had no clear phenotypic benefit from DCA, although a metabolic improvement was seen ^12^. We postulate that this may relate to DCA requiring a certain minimum expression of PDHc to have therapeutic benefit, as it activates residual PDHc and in the setting of severe DLD knockout would anticipate insufficient substrate present for meaningful activation of residual PDHc activity by DCA. Consistent with this hypothesis, studies in the *C. elegans dld-1* models presented here revealed significant therapeutic benefit of DCA on survival, neuromuscular and metabolic phenotypes primarily in the moderate (1:20) and/or mild (1:100) *dld-1* dilution strains, rather than the full-dose *dld-1(RNAi)*, although linear animal growth and neuromuscular activity was significantly improved in all three RNAi strains studied upon DCA treatment.

Collectively, these preclinical evaluations demonstrate objective therapeutic benefit of DCA in two distinct genetic etiologies of PDHc deficiency, namely homologues of *PDHA1* and *DLD*. Additional insight is provided into the selective sensitivity of PDHc deficiency models to DCA, showing that beneficial effects in rescuing mitochondrial stress that is significantly induced in the *dld-1 (RNAi)* models inversely correlated with disease severity. Importantly, these preclinical data serve as confirmatory evidence to support data from a single phase 3 clinical trial completed to evaluate the therapeutic utility of DCA in PDHc deficiency, with demonstrated beneficial effects on survival, healthspan, tissue lactate, and diverse aspects of mitochondrial physiology. While that clinical trial excluded *DLD* deficiency patients due to the multi-enzyme deficiency caused by the E3 subunit being a component of multiple mitochondrial keto acid dehydrogenases beyond PDHc, these preclinical data in *dld-1(RNAi)* knockdown worms suggest the survival, health span, as well as diverse metabolic disruptions that occur in mild to moderate degrees of *DLD* deficiency do significantly improve with DCA treatment.

## AUTHOR CONTRIBUTIONS

MJF conceived of the study and obtained funding and oversaw study performance. CR performed *C. elegans* growth, sequencing, phenotyping, and treatment studies. VM, CR and NM performed *C. elegans* lifespan studies. CR and VM performed the western blot analysis. TO performed respiration analyses studies. SH performed the confocal imaging experiments and assisted with analysis of both Seahorse and confocal microscopy data. ML performed confocal microscopy data analyses. KK analyzed the automatic worm locomotor activity data. ENO performed lactate and pyruvate analyses. RX provided statistical review and guidance. CR and VEA analyzed the data, performed statistical analyses, and wrote the initial manuscript draft with MJF. All authors reviewed and approved the final manuscript version.

## ACKNOWLEDGEMENTS

This study was funded in part by a researcher initiated sponsored research agreement with Saol Therapeutics and the National Institutes of Health (R35-GM134863, M.J.F., PID). Dichloroacetate studied in this work was provided under sponsored research agreement from Saol Therapeutics. The content is solely the responsibility of the authors and does not necessarily represent the official views of the NIH.

## ETHICS APPROVAL AND PATIENT CONSENT STATEMENT

No ethics approval was required nor obtained for the work performed in this study, which was limited to analyses in the invertebrate animal *C. elegans.* No vertebrate animals, patient samples, or patient data were included in this study.

## DATA SHARING STATEMENT

The data that support the findings of this study are available in the supplementary material of this article.

## COMPETING INTERESTS

MJF is inventor of US patent 12,011,452 B2 issued Jun 18, 2024, “Compositions and Methods for Treatment of Mitochondrial Respiratory Chain Dysfunction and Other Mitochondrial Disorders”. MJF is engaged with companies involved in mitochondrial disease therapeutic preclinical and/or clinical-stage development. MJF is co-founder of Rarefy Therapeutics; an advisory board member with equity interest in RiboNova Inc.; a scientific advisory board member and paid consultant with Khondrion and Larimar Therapeutics; has been a paid consultant for Ajinomoto-Cambrooke, Inc., Astellas (formerly Mitobridge), BPGbio, Casma Therapeutics, Cyclerion Therapeutics, Epirium Bio (formerly Cardero Therapeutics), HealthCap VII Advisor AB, Imel Therapeutics, Mayflower, Inc., Primera Therapeutics, Inc., Minovia Therapeutics, Mission Therapeutics, NeuroVive Pharmaceutical AB, Precision Biosciences, Reneo Therapeutics, Saol Therapeutics, Sironax, Stealth BioTherapeutics, Vincere Bio, and Zogenix; and/or has been a sponsored research collaborator for Aadi Bioscience, Adjuvia Therapeutics, Astellas, Cyclerion Therapeutics, Epirium Bio, Imel Therapeutics, Khondrion, Merck, Minovia Therapeutics, Mission Therapeutics, NeuroVive Pharmaceutical AB, Precision Biosciences, Pretzel Therapeutics, PTC Therapeutics, Raptor Pharmaceuticals, REATA Inc., Reneo Therapeutics, RiboNova, Saol Therapeutics, Standigm, Stealth BioTherapeutics, and Thiogenesis. MJF also has received royalties from Elsevier and speaker fees from Agios Pharmaceuticals, GenoMind, and educational honoraria from PlatformQ and Chemistry Rx. No other co-author ha**s a** relevant conflict of interest to declare.

Copyright © 1993-2023, University of Washington, Seattle. GeneReviews is a registered trademark of the University of Washington, Seattle. All rights reserved.).

