## supplemental files for "Dichloroacetate improves animal survival, growth, neuromuscular activity, mitochondrial stress and physiology, and elevated lactate in *C. elegans pdha-1* and *dld-1* RNAi models of pyruvate dehydrogenase complex deficiency (PDCD)"

**Supplementary Fig. 1. (A)** Amino acid alignment between the human PDHA1and *C. elegans* PDHA-1 proteins. **(B)** Uncropped western blot images corresponding to Fig. 1A, together with two additional biological replicates used for PDHA-1 protein quantification in Fig. 1B. **(C)** Lifespan analysis of wild-type worms (N2 Bristol) treated with a 25 mM DCA shows no significant differences compared with untreated controls, measured by the WormScan assay, in presence of FUDR. **(D)** Manual lifespan analysis without FUDR showed significant reduction in lifespan upon *pdha-1(RNAi)* knockdown relative to the wild-type (N2 Bristol) control (p=0.0012), whereas treatment with 25 mM DCA restored lifespan in *pdha-1(RNAi)* knockdown worms toward wild-type (N2 Bristol) levels (p=0.013), across three biological replicates. **(E)** Stimulated locomotor activity of wild-type (N2 Bristol) and *pdha-1(RNAi)* worms measured using an automated system, under untreated conditions and across a four-point DCA dose series. Analyses are shown for stages L4+2 days, L4+5 days, and L4+10 days of adulthood. Data represent the mean ± SD of 6 biological replicates with ~20 animals each. (**F**) Log-linear regression analysis of DCA dose–response on worm length, demonstrating a significant positive dose–response relationship between DCA concentration and normalized worm length.

**Supplementary Fig. 2.** Microscopy images of wild-type (N2 Bristol) (A) and *pdha-1(RNAi)* knockdown (B) worms treated with water (buffer control) or a four-point DCA dose series, imaged at stage L4+1 day. Scale bar: 200 µm

**Supplementary Fig. 3.** Microscopy images of wild-type (N2 Bristol) (A) and *pdha-1(RNAi)* knockdown (B) worms treated with water (buffer control) or a four-point DCA dose series, imaged at stage L4+5 day. Scale bar: 200 µm

**Supplementary Fig. 4. (A)** Representative CX5 images for worms fed with empty vector (wild-type, *hsp-6p::GFP; myo-2p::mCherry*) and *pdha-1*(*RNAi*) bacteria, treated with water or 25 mM DCA. The red fluorescence (mCherry) was used to determine the number of animals per well, and the green fluorescence (GFP) was used to quantify the level of $UPR^{mt}$. **(B-C)** FCCP titration curve was performed untreated wild-type (N2 Bristol) (B) and untreated *pdha-1(RNAi)* (C) animals to determine the optimal FCCP concentration for Seahorse analyses. **(D)** Oxygen consumption rate (OCR) of wild-type (N2 Bristol) and *pdha-1(RNAi)* knockdown animals treated with water (buffer control) or 25 mM DCA, measured by

Seahorse assay following sequential addition of two mitochondrial function modulators: FCCP (25 μM) and sodium azide (50 mM). Data represent the mean ± SD of 15 technical replicates per biological replicate (with ~75 animals/well). **(E)** Non-mitochondrial OCR measured for the wild-type (N2 Bristol), *pdha-1(RNAi)*, and *dld-1(RNAi)* worms, treated with water or 25 mM DCA, measured across 3 biological replicates. Data represent the mean ± SD of 3 replicates with 15 technical replicates each.

**Supplementary Fig. 5.** Representative Biosorter dot plots showing time of flight (TOF) versus green fluorescence (left panels) and red fluorescence (right panels) for wild-type worms (N2 Bristol) **(A)**, *pdha-1(RNAi)* worms **(B)**, and *pdha-1(RNAi)* worms treated with 25 mM DCA **(C)**. Each dot represents an individual animal. All objects detected were included in the analysis; no gating was applied. Panels shown are representative of three independent biological replicate experiments per condition.

**Supplementary Fig. 6. (A-C)** CX5 analysis of wild-type (*hsp-6p::GFP; myo-2p::mCherry*) and *pdha-1(RNAi)* knockdown worms treated with water or 25 mM DCA and co-labeled with TMRE and MTG. Average red **(B)** and green **(C)** fluorescence of *pdha-1(RNAi)* animals are shown, normalized to wild-type levels. Data represent the mean ± SD of 4 replicates with ~50 animals each. **(D)** CX5 analysis of the relative mitochondrial membrane potential, calculated as the ratio of TMRE (red channel) fluorescence to MTG (green channel) fluorescence, relative to wild-type (N2 Bristol). Data represent the mean ± SD of 4 replicates with ~50 animals each. **(E)** Pyruvate/Lactate ratio in the *pdha-1(RNAi)* knockdown worms was increased relative to wild-type (N2 Bristol), while treatment with DCA did not significantly alter this ratio. *: $p < 0.05$ **(F)** Representative confocal microscopy images of wild type (*myo-3p::GFP(mit)*), *pdha-1* and *dld-1(RNAi)* animals treated with water or 25 mM DCA. Scale bar: 10 µm

**Supplementary Fig. 7. (A)** Amino acid alignment between the human DLDH and *C. elegans* DLD-1 protein. **(B)** Uncropped western blot images corresponding to Fig. 3A, together with two additional biological replicates used for quantification of the DLD-1 protein in Fig. 3B.

**Supplementary Fig. 8.** Microscopy images of *dld-1, dld-1 (1:20) and dld-1 (1:100) RNAi* worms treated with water (buffer control) or a four-point DCA dose series, imaged at the stage L4+1 day. Scale bar: 200 μm

**Supplementary Fig. 9.** Microscopy images of *dld-1, dld-1 (1:20) and dld-1 (1:100) RNAi* worms treated with water (buffer control) or a four-point DCA dose series, imaged at stage L4+5 days. Scale bar: 200 μm

**Supplementary Fig. 10. (A)** Manual lifespan analysis without FUDR showed that *dld-1(1:20 RNAi)* worms had significantly reduced survival compared to wild-type controls (N2 Bristol, $p<0.0001$), whereas treatment with 25 mM DCA significantly increased their survival ($p<0.0001$) across three biological replicates. **(B)** Lifespan analysis using the WormScan method in presence of FUDR showed that *dld-1(1:100 RNAi)* worms had significantly reduced survival compared to wild-type controls (N2 Bristol, $p=0.002$), whereas treatment with 25 mM DCA had no effect on survival. Three biological replicates are presented. **(C)** Lifespan analysis using a standard manual assay without FUDR showed no difference in survival of *dld-1(1:100 RNAi)* knockdown worms compared to wild-type controls (N2 Bristol), and treatment with 25 mM DCA did not affect survival. Three biological replicates are presented. **(D)** Lifespan analysis using a standard manual assay without FUDR at 20ºC showed that *dld-1* knockdown worms had significantly increased survival compared to wild-type controls (N2 Bristol, $p<0.0001$), while treatment with 25 mM DCA further significantly extended their survival ($p=0.01$). Three biological replicates are presented. **(E)** Light stimulated locomotor activity of wild-type (N2 Bristol, WT) and *dld-1* worms measured using an automated system under untreated conditions and across a four-point DCA dose series. Analyses are shown for the L4+2 days, L4+5 days, and L4+10 days of adulthood. Data represent the mean ± SD of 6 technical replicates with ~20 animals each.

**Supplementary Fig. 11. (A)** Representative CX5 images for worms fed with L4440 (*hsp-6p::GFP* + *myo-2p::mCherry*, wild type) and *dld-1(RNAi)*, *dld-1(1:20 RNAi)*, and *dld-1(1:100 RNAi)* bacteria treated with water or 25 mM DCA. The red fluorescence (mCherry) was used to determine the number of animals per well, and the green fluorescence (GFP) was used to quantify the level of mitochondrial stress induction. **(B)** Quantification of mitochondrial stress upon feeding worms with increasing concentrations of *dld-1* RNAi bacteria. Data represents the mean ± SD of 8 replicate experiments with ~50 animals per condition. **(C)** An FCCP titration curve was performed in untreated *dld-1(RNAi)* knockdown animals to determine the optimal FCCP concentration for Seahorse analyses. Data represent the mean ± SD of 15 technical replicates per biological replicate, with one animal per condition. **(D)** OCR of wild-type (N2 Bristol) and *dld-1(RNAi)* knockdown animals treated with water (buffer control) or 25 mM DCA, measured by Seahorse assay following sequential addition of mitochondrial function modulators: FCCP (25 µM) and sodium azide (50 mM). Data represent the mean ± SD of 15 technical replicates per biological replicate, with one animal per condition. **(E)** Pyruvate/Lactate ratio in *dld-1*(*1:20* RNAi) knockdown worms trended toward increase relative to wild type (N2 Bristol), and treatment with 25 mM DCA trended to further increase this ratio. *dld-1(1:100 RNAi*) worms showed no difference in pyruvate/lactate levels compared to wild type, and treatment with DCA did not significantly change this ratio. ****: $p<0.0001$

**Supplementary Fig. 1.**

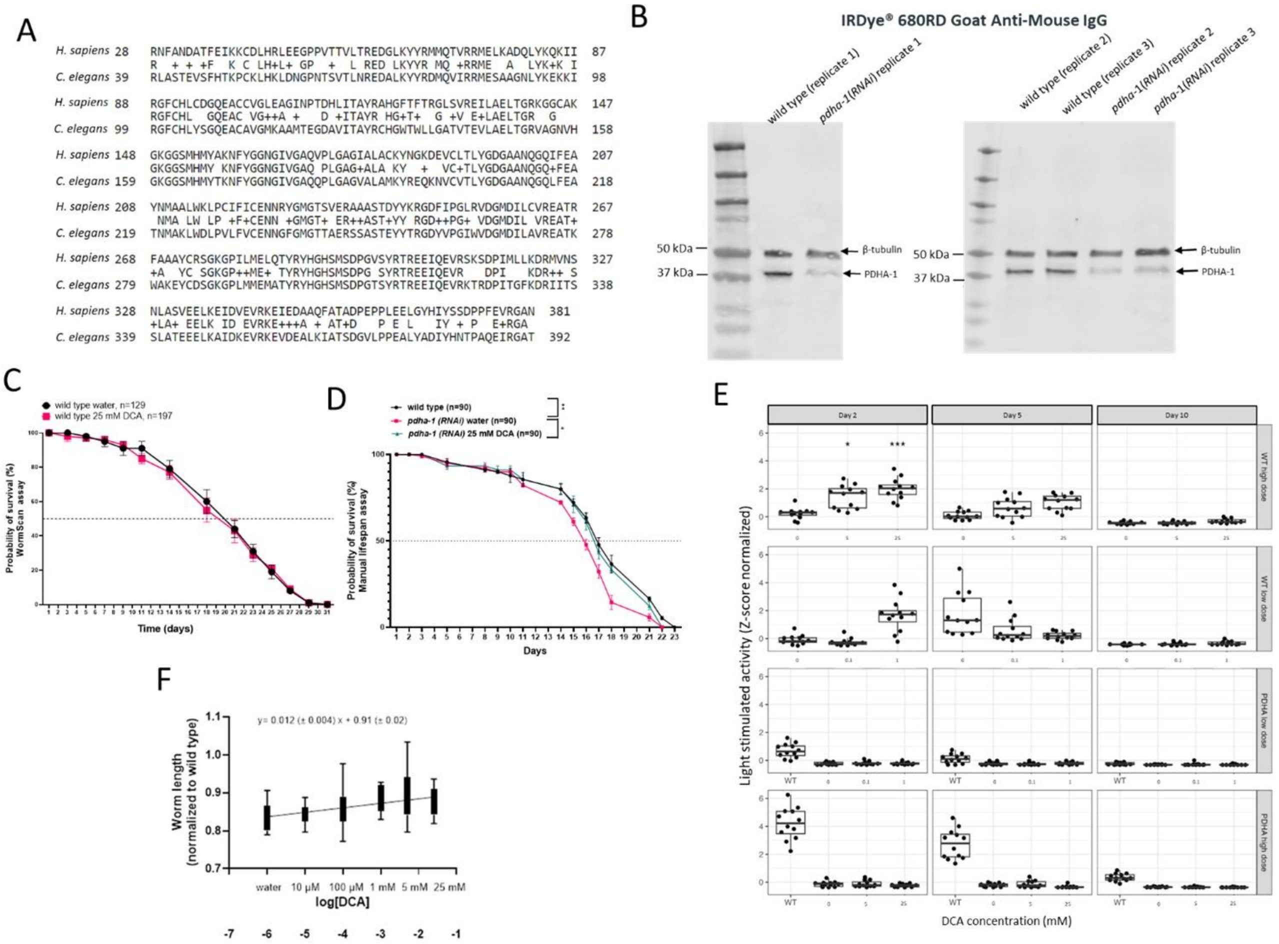


**Supplementary Fig. 2.**

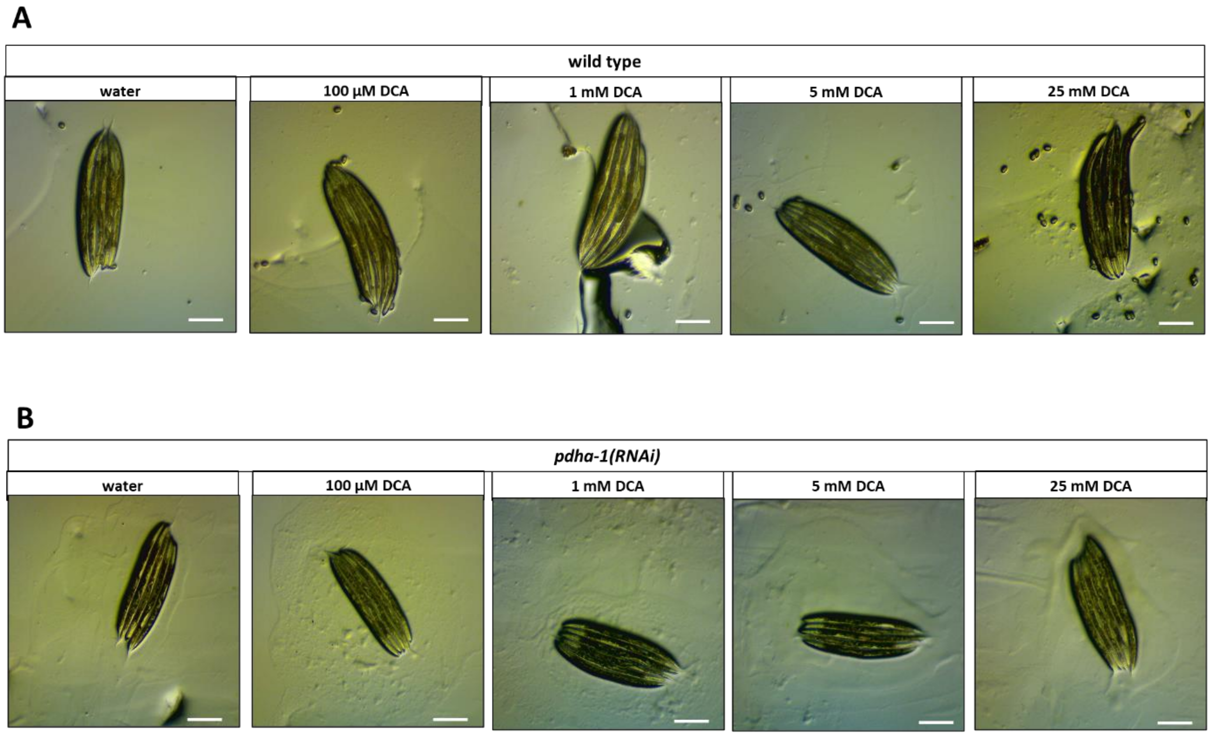


**Supplementary Fig. 3**

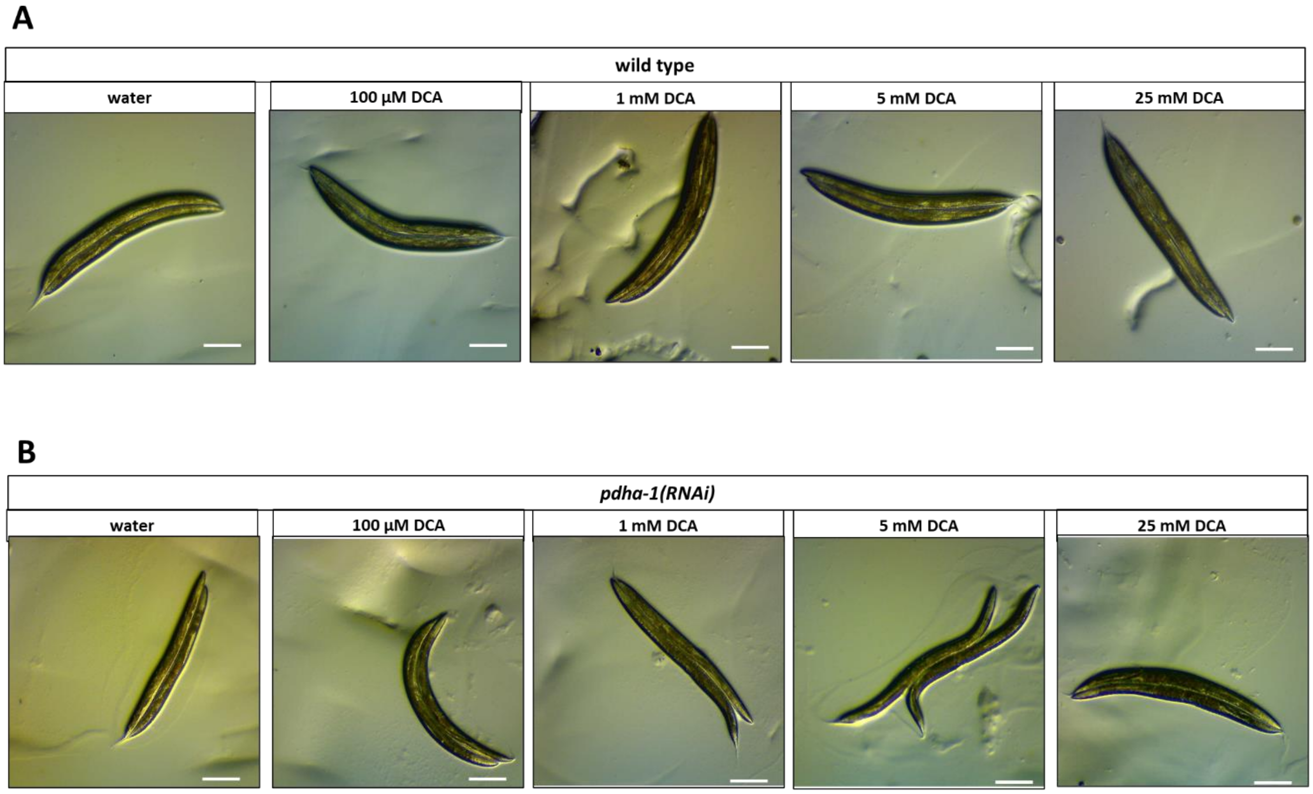
A
wild type
water
100 µM DCA
1 mM DCA
5 mM DCA
25 mM DCA
B
pdha-1(RNAi)
water
100 µM DCA
1 mM DCA
5 mM DCA
25 mM DCA

**Supplementary Fig. 4.**

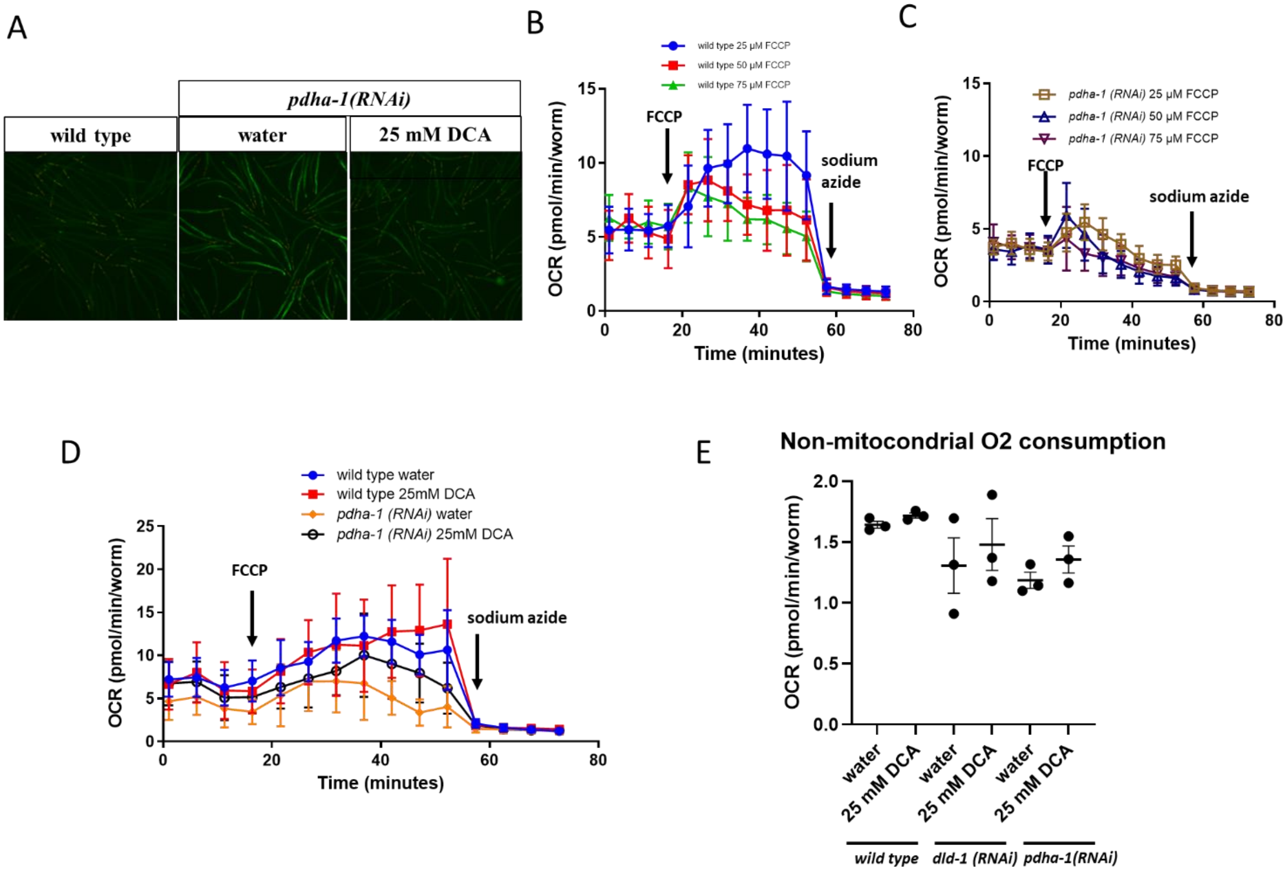

**Supplementary Fig. 5.**

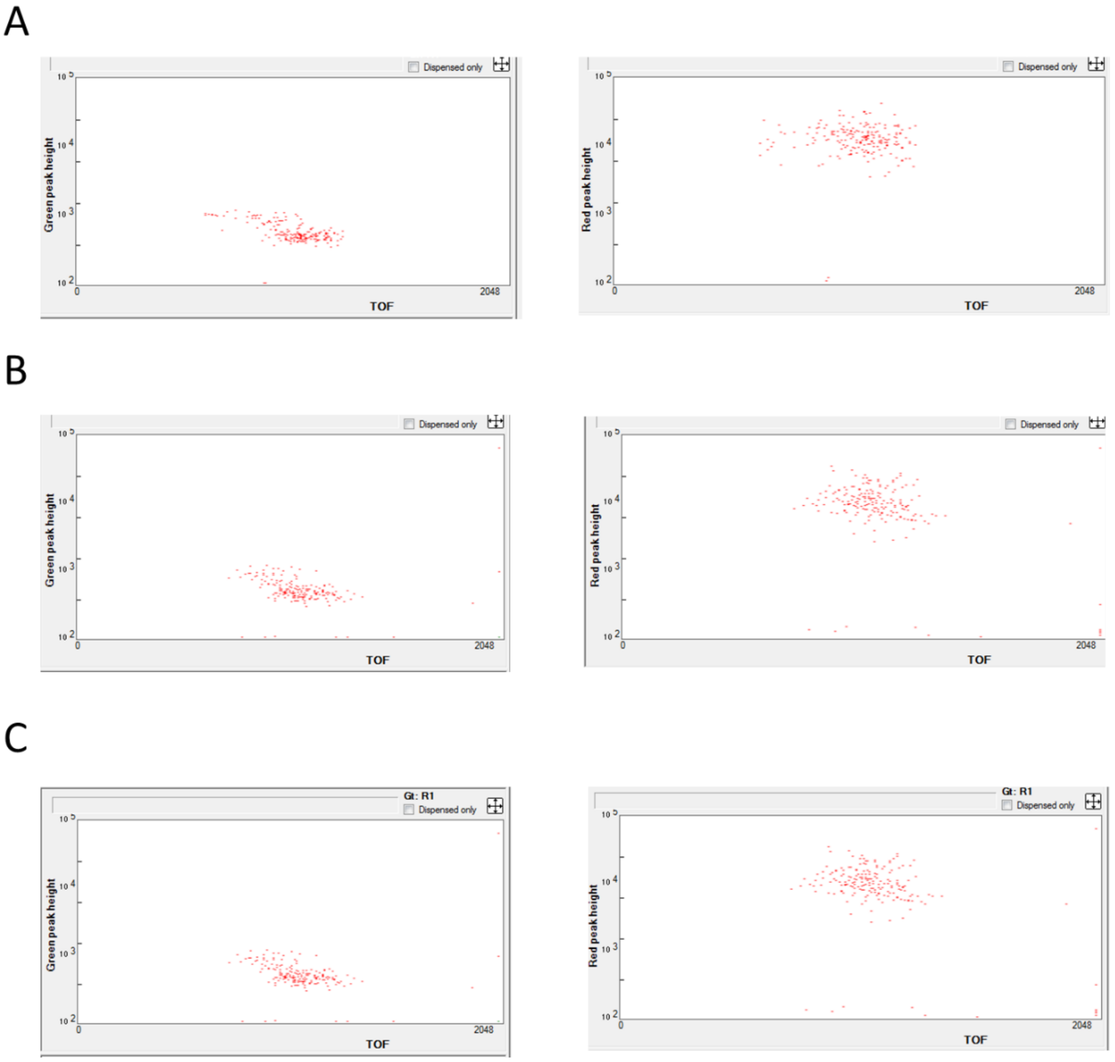

**Supplementary Fig. 6.**

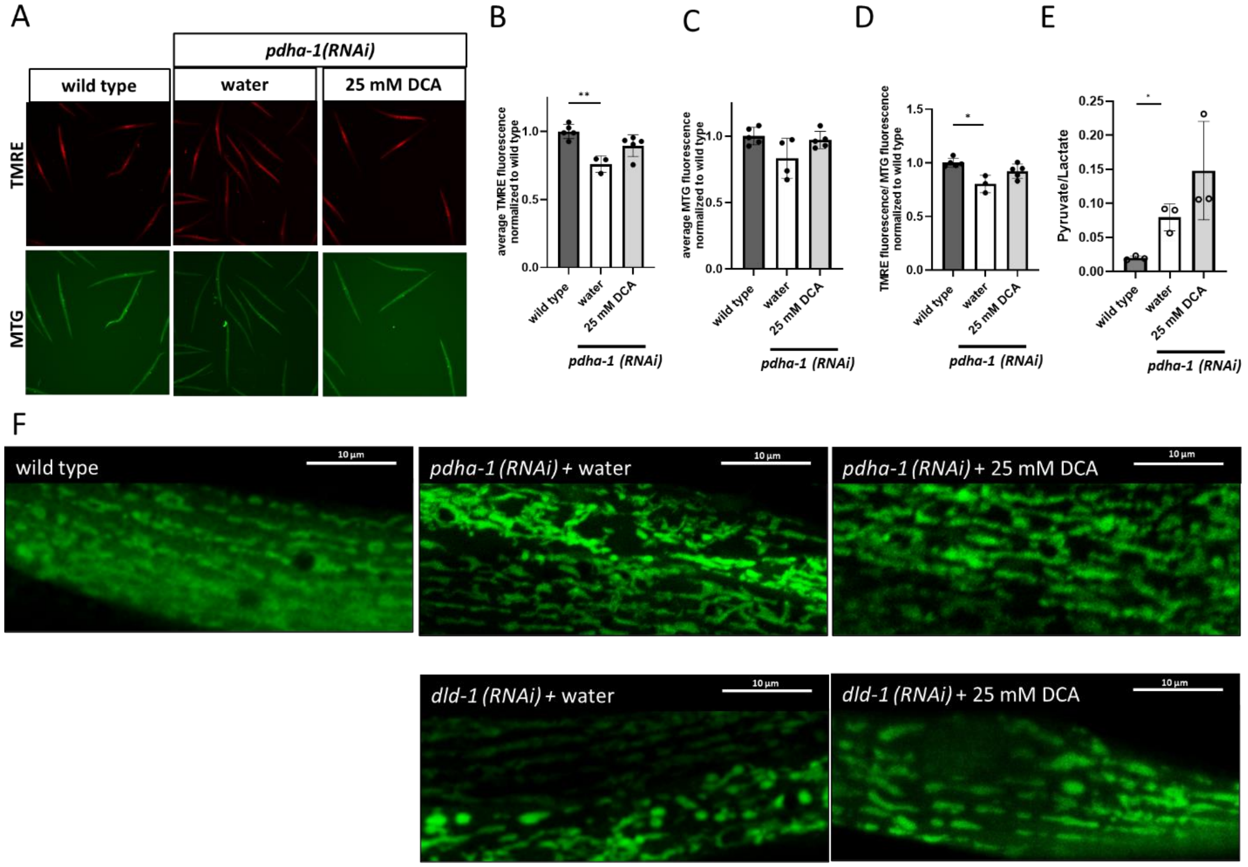

**Supplementary Fig. 7.**

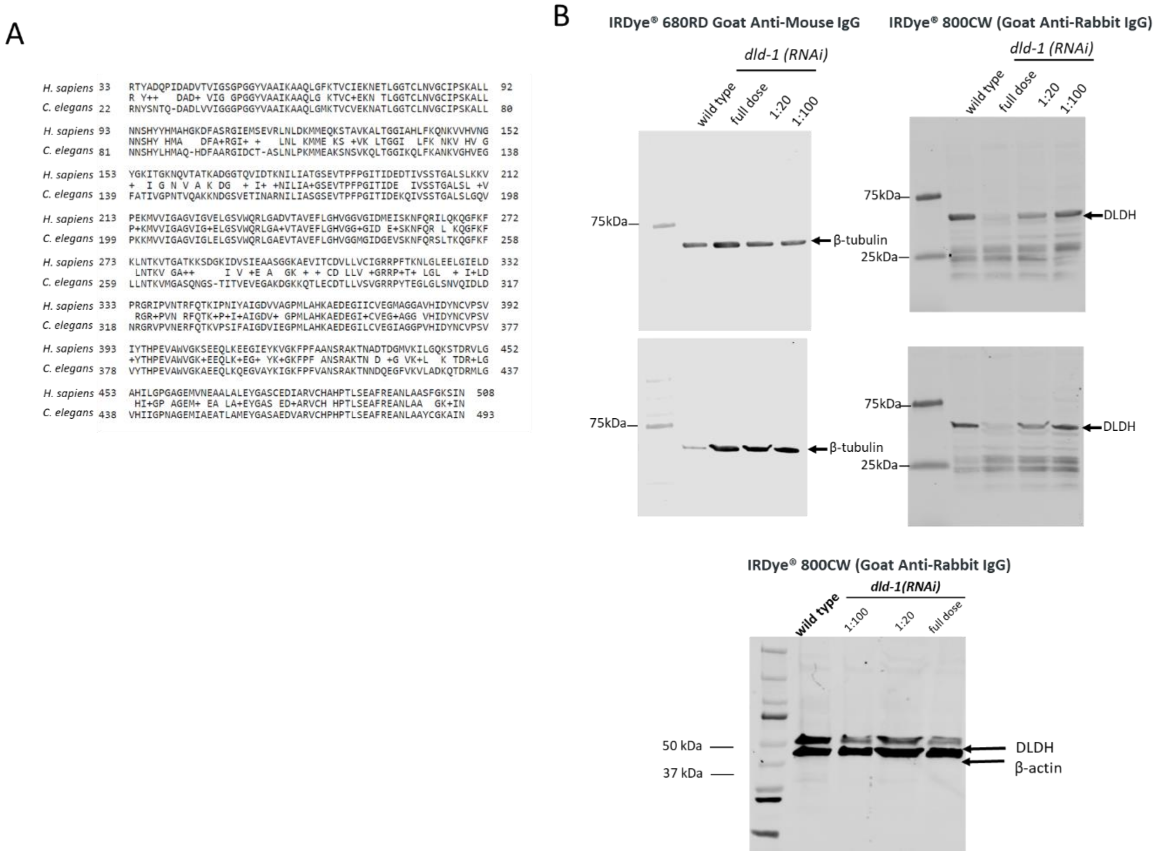

### Supplementary Fig. 8.

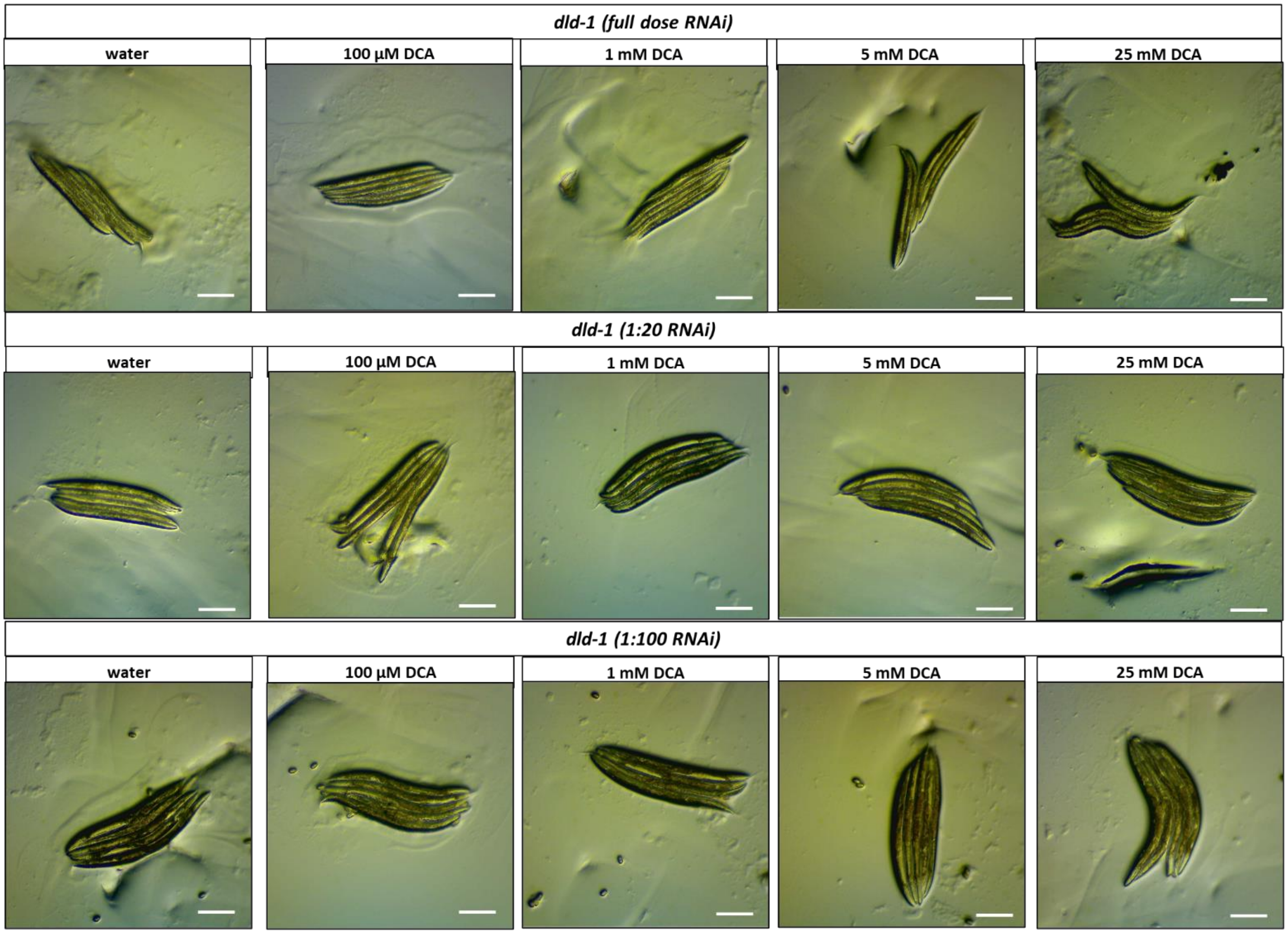

### Supplementary Fig. 9.

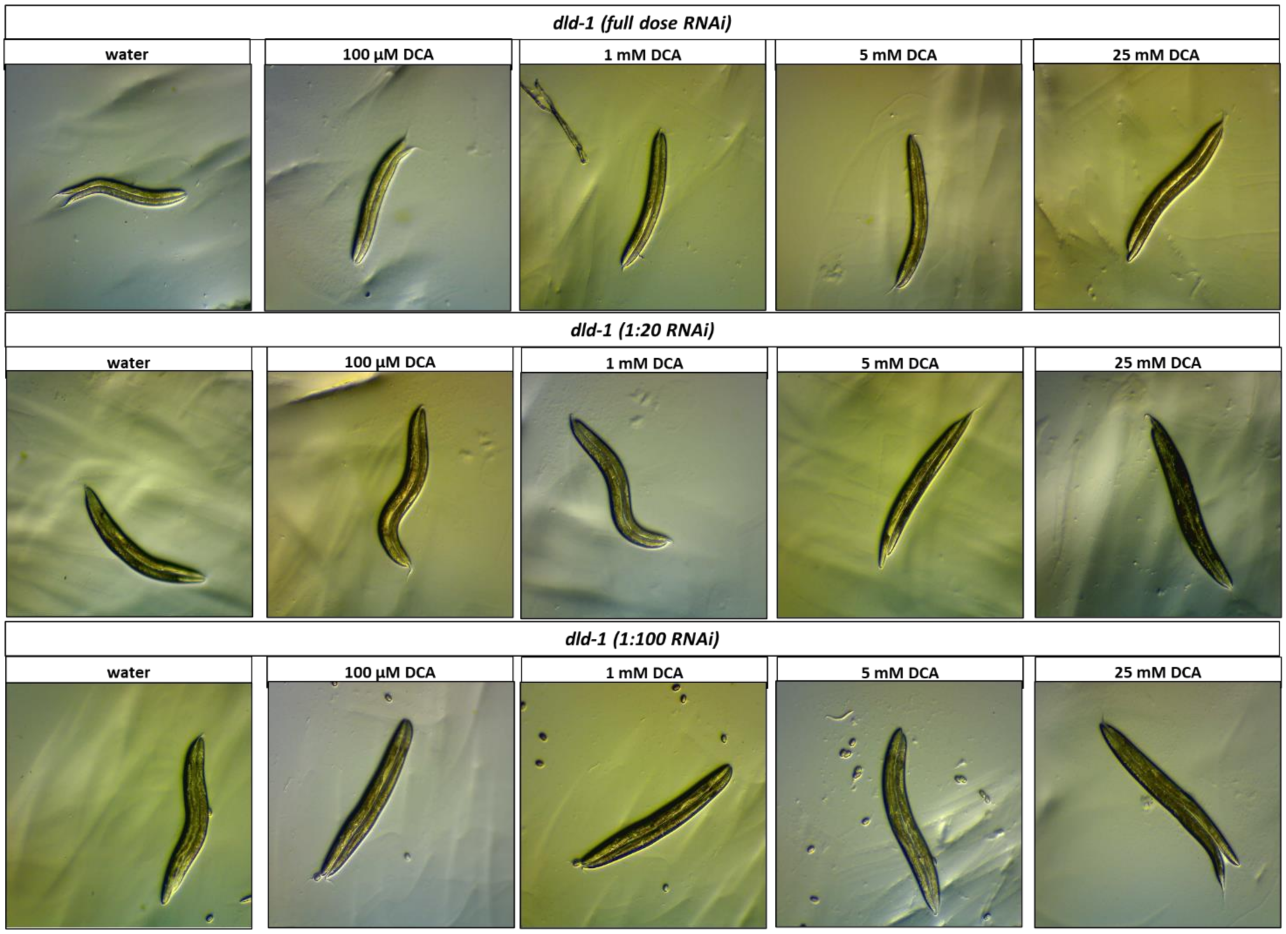

**Supplementary Fig. 10.**

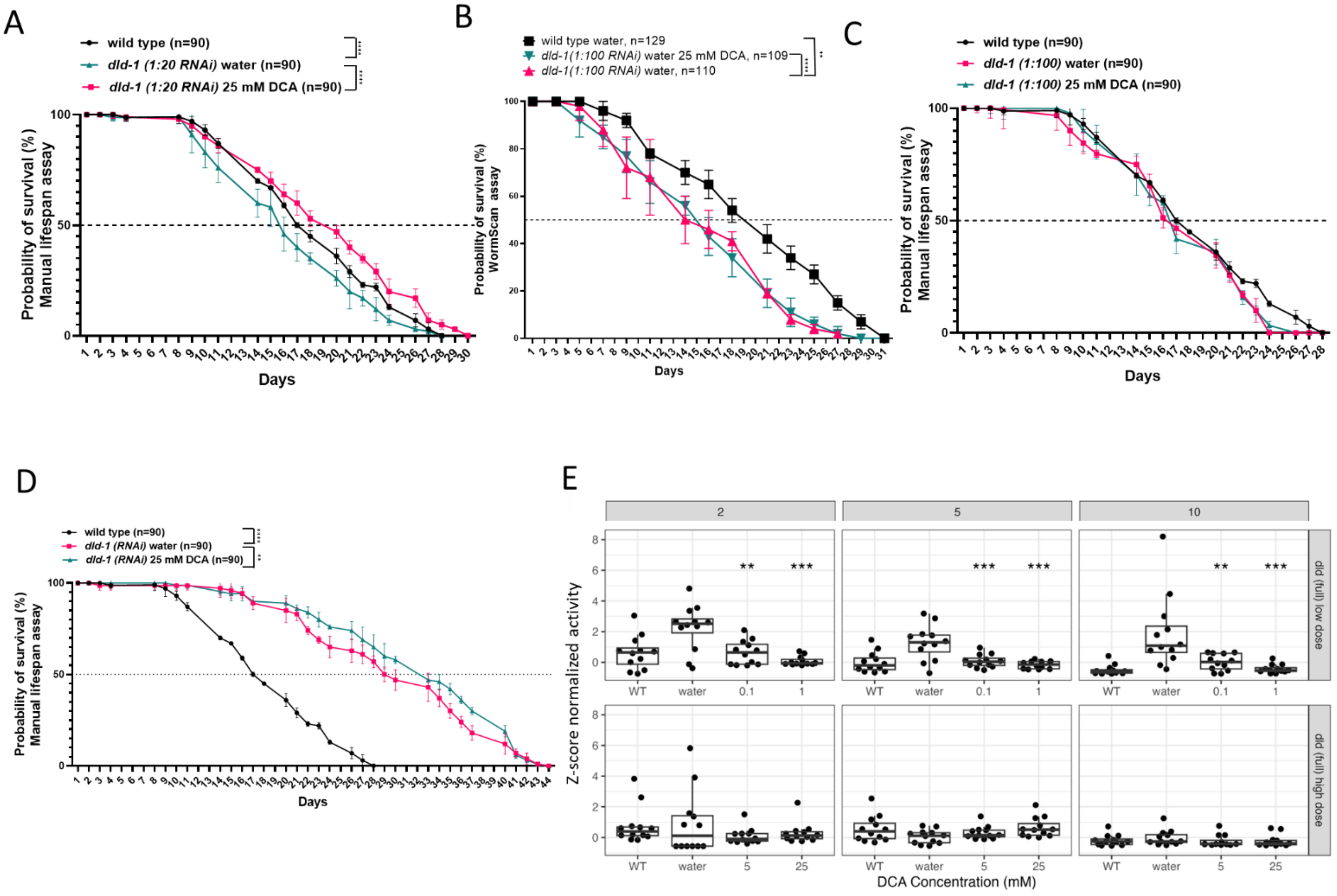

**Supplementary Fig. 11.**

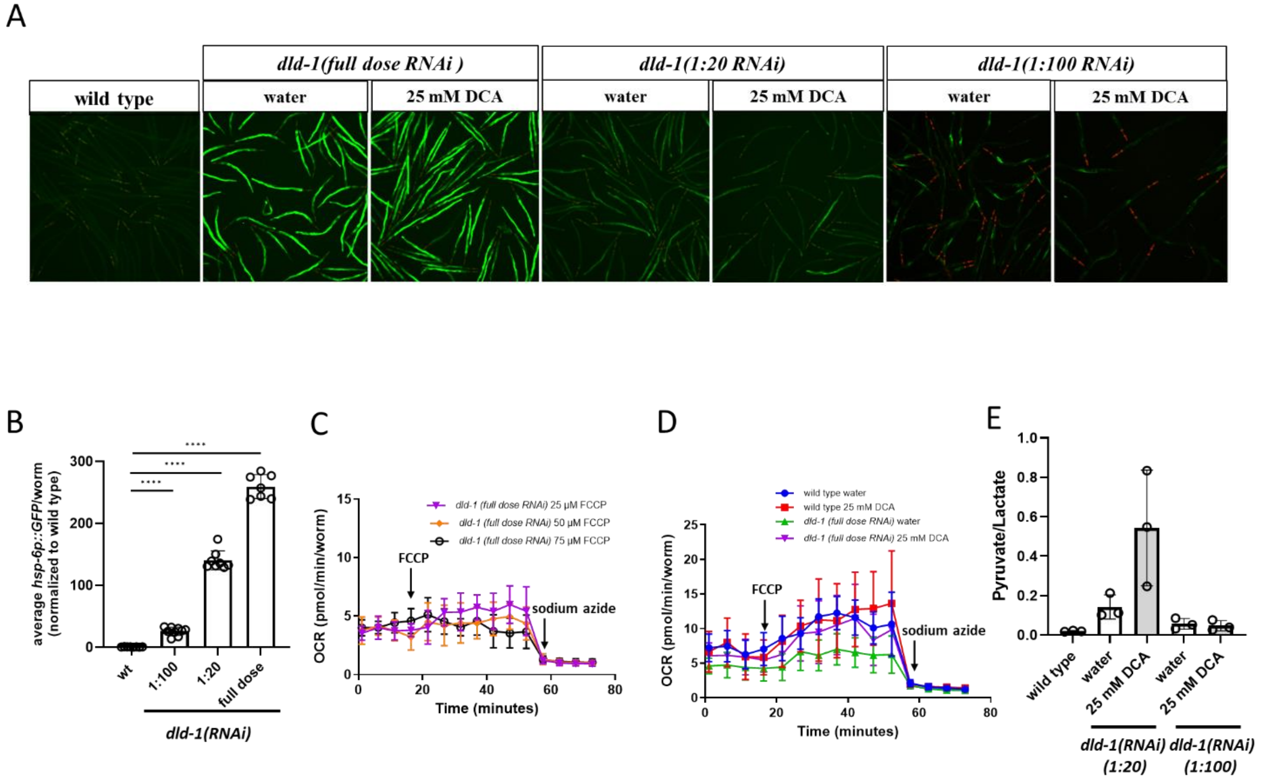

**Supplementary Table 1.** Comparison of lifespan data, measured using the WormScan method, in presence of FUDR, for *dld-1(RNAi)* knockdown worms from this study and previous reports.

| | ***wild-type*** | | ***dld-1(RNAi)*** | | ***dld-1 (1:20 RNAi)*** | | ***dld-1(1:100 RNAi)*** | ***pdha-1(RNAi)*** |
|---|---|---|---|---|---|---|---|---|
| | previous study[1] (N2 Bristol) | current study (N2 Bristol) | previous study[1] | current study | previous study[1] | current study | current study | current study |
| **Level of protein knockdown determined by western immuno-blotting (relative to wild type)** | N/A | N/A | 71% | 80% | 38% | 56% | 18% | 70% |
| **DCA treatment onset** | L4 | embryo | L4 | embryo | L4 | embryo | embryo | embryo |
| **Median lifespan (days, in water control)** | 21 | 21 | 28 | 14 | 19 | 16 | 16 | 12 |
| **Median lifespan (days, in 25 mM DCA treatment)** | not assessed | 20 | 32 | 19 | 25 | 20 | 16 | 13 |

**Supplementary Table 2.** Overview of the experimental design and conditions presented in this study

| **Experiment** | **Strain** | **Treatment start** | **Measurement time point** |
|---|---|---|---|
| Lifespan (manual and WormScan assays) | N2 Bristol | embryo | Imaged from stage L4, for 30 days |
| Stimulated locomotor activity measurements (automated light-stimulated activity) | N2 Bristol | embryo | Imaged from stage L4 for 30 days |
| Worm linear growth | N2 Bristol | embryo | L4+1 day |
| Neuromuscular activity (thrashing) | N2 Bristol | embryo | L4+1 day |
| $UPR^{mt}$ stress (high-content imaging) | *hsp-6p::GFP; myo-2p::mCherry* | embryo | L4+1 day |
| Oxygen consumption rate (seahorse analysis) | N2 Bristol | embryo | L4 |

| | | | |
|---|---|---|---|
| Mitochondrial membrane potential, mitochondrial mass (Biosorter analysis) | N2 Bristol | embryo | L4+1 day (fluorescent labeling started at stage L4) |
| Measurement of lactate and pyruvate levels | N2 Bristol | embryo | L4+1 day |
| Mitochondrial morphology (confocal microscopy) | SJ4103 strain carrying the zcls14 construct (*myo-3p::GFP(mit)*) | embryo | L4+1 day |
